# Systematic analysis of Epstein-Barr virus genes and their individual contribution to virus production and composition

**DOI:** 10.1101/2021.04.26.439973

**Authors:** Yen-Fu Adam Chen, Bianca Mocanu, Ezgi Akidil, Dagmar Pich, Josef Mautner, Wolfgang Hammerschmidt

## Abstract

Epstein-Barr virus (EBV), a member of the large herpesvirus family is a very complex human *γ*-herpesvirus. Its complexity with respect to viral gene regulation, its preferred latent life style and technical difficulties are major obstacles, which hinder efficient virus generation *in vitro* to mass-produce or establish recombinant wild-type EBV or mutant derivatives. To explore conditions optimizing and improving virus production, we established and tested an EBV gene library with 77 expression plasmids and a set of designed shRNAs. With this tool set we investigated the contributions of individual viral genes in the context of virus synthesis and virion functions. Engineered virus stocks were systematically analyzed with respect to physical and bioparticle concentration, virus titer and virus uptake by primary human B cells, EBV’s target cells *in vivo*. To quantitate virus uptake by these cells, we developed a novel *ß*-lactamase-based assay that can monitor fusion events of the viral envelope with membranes of recipient cells at the level of single cells. Based on our findings, EBV does not encode a dominant regulator that governs virus production, but our results identified several EBV genes such as BALF4, BVLF1 and BKRF4, encoding a viral glycoprotein, a transcriptional regulator of late viral genes, and a possible tegument protein, respectively, that improve virus production regarding virus yield, virus composition and quality, and virus uptake by primary human B cells.

**Importance:** For more than 20 years HEK293 cells have been instrumental to produce virus stocks of recombinant EBV. The identification of this cell line as a source of infectious EB virions was unexpected because HEK293 cells are very distant from B cells and in particular plasma cells which produce EBV progeny *in vivo*. To our knowledge, no systematic analysis has addressed the fundamental question whether virus yield with respect to virus concentration, virion composition and functionality can be improved to produce recombinant EBV stocks in HEK293 cells. We tackled this question and analyzed 77 individual EBV genes for their possible contribution to virus yield. To analyze and compare different virus stocks we used established and developed novel assays to characterize important parameters of EB virions and their functions.

## Introduction

Epstein-Barr virus (EBV), also known as human herpesvirus 4 (HHV4) (Albà et al., 2001) was discovered by Michael Anthony Epstein, Bert Geoffrey Achong and Yvonne Barr in samples of Burkitt’s lymphoma biopsies in 1964 (Epstein et al., 1964). EBV together with Kaposi sarcoma-associated herpesvirus (KSHV, HHV-8) are two human herpesviruses that belong to the gamma herpesvirus subfamily. A hallmark of herpesviruses is their ‘lifestyle’ characterized by predominantly latent infections *in vivo*. Latency means that the infected target cells carry viral genome copies, which are epigenetically repressed such that no or only a very restricted set of viral genes is expressed preventing immune recognition as well as virus *de novo* synthesis. Only upon reactivation, the latently infected cells turn into virus factories which release progeny. EBV adopts this lifestyle also *in vitro* and thus is a prime model of herpesvirus latency.

*in vivo* and *in vitro*, EBV very efficiently infects mature human B cells to establish a latent infection in them. *in vitro*, EBV reprograms the infected B cells in the pre-latent phase of infection (Mrozek-Gorska et al., 2019) and turns the formerly quiescent B cells into proliferating lymphoblasts within only a few days, which emerge as stable lymphoblastoid cell lines (LCLs). The majority of the cells maintains the latent state during *in vitro* propagation whereas a small, variable fraction spontaneously enters EBV’s lytic, productive phase and releases infectious virions. LCLs that support the latent phase express a very restricted viral program which includes few latent EBV genes, only. Very much in contrast, cells that support virus production express the entire set of lytic viral genes. A single viral gene called BZLF1 (also termed ZEBRA, Zta, Z or EB1) is essential and capable of initiating EBV’s lytic phase in latently infected cells (Feederle et al., 2000). It is unclear what triggers BZLF1’s expression in LCLs *in vitro* but when latently EBV-infected B cells encounter their cognate antigen *in vivo* they can differentiate into pre-plasmablasts and plasma cells. Upon terminal differentiation of these B cells, BZLF1 is expressed, which initiates escape from latency and the downstream expression of all lytic viral genes leading to *de novo* synthesis of viral progeny (Laichalk and Thorley-Lawson, 2005).

BZLF1 is a homodimeric bZIP transcription and pioneer factor (Schaeffner et al., 2019) and binds two classes of BZLF1 (or ZEBRA) response elements (ZREs): CpG-free motifs resembling the consensus AP-1 site recognized by cellular bZIP proteins such as c-Jun/c-Fos (Farrell et al., 1989), and CpG-containing motifs that are selectively bound by BZLF1 upon cytosine methylation (Bhende et al., 2004; Dickerson et al., 2009; Bergbauer et al., 2010; Flower et al., 2011; Woellmer et al., 2012; Tillo et al., 2017; Buschle et al., 2021; Bernaudat et al., 2021). EBV’s virion DNA, which is unmethylated upon infection (Kalla et al., 2010), precludes the binding of BZLF1 to certain promoters of essential lytic viral genes in the pre-latent phase of infection. The requirement of CpG methylated viral DNA to support EBV’s lytic phase also means that EBV cannot turn a newly infected cell into a viral factory (Kalla et al., 2010; Woellmer et al., 2012; Buschle et al., 2021). Instead, EBV establishes a strictly latent infection initially and escapes from it later when viral DNA has reached a certain level of CpG methylation (Kalla et al., 2010, 2012). In this way, the EBV virus has created its own time-dependent, epigenetic switch to control its biphasic life cycle (Buschle and Hammerschmidt, 2020).

EBV’s biphasic mode has been an obstacle to study this virus genetically as it lacks a tractable lytic cycle in which to generate and from which to select viral mutants. This is contrary to herpesviruses of the alpha and beta subfamilies, which infect cells to give rise to progeny readily and within a few days. As a consequence, genetic manipulation of EBV’s genomic DNA has not been possible for decades after its discovery in the sixties very much in contrast to herpes simplex virus 1. This virus’ era of molecular genetics started more than 40 years ago in the Roizman lab using forward selection in cells that support lytic viral replication (Post et al., 1980; Mocarski et al., 1980; Post and Roizman, 1981). The beginning of EBV’s genetic era dates back to 1989, when viral vector techniques and cells infected with EBV variants showed that this virus is also amendable to targeted genetic analysis in principle (Hammerschmidt and Sugden, 1989). Alternatively, Cohen et al. used a similar approach based on homologous recombination (Cohen et al., 1989). Although this technology was a pioneering step in the EBV field (Kempkes et al., 1995c), its practical application was difficult, cumbersome and limited as pointed out in a review by Feederle et al. (Feederle et al., 2010).

Reverse molecular genetics of EBV’s latent genes became more versatile only after the artificial assembly of half-sized EBV genomes in E. coli. They were established with the aid of single copy mini-F-factor plasmids and large fragments (O’Connor et al., 1989) of molecularly cloned virion DNA. The so-called mini-EBV genomes were functional and transformed human primary B-lymphocytes after their direct or indirect delivery and provided direct genetic access to all latent genes of EBV (Kempkes et al., 1995b, a; Kilger et al., 1998).

The eventual breakthrough came when we introduced the same single copy mini-F-factor plasmid into LCLs infected with the B95.8 lab strain of EBV to provoke targeted homologous recombination of mini-F-factor and genomic EBV DNAs in these cells (Delecluse et al., 1998). Only then it was possible to establish the entire EBV genome in E. coli as a bacterial artificial chromosome, a BACmid, which we also call maxi-EBV. In E. coli, the maxi-EBV DNA is accessible to (almost) unlimited genetic manipulations because the large EBV insert is non-functional in the prokaryotic host. Genetic alterations in E. coli also follow the principle of homologous DNA recombination, which is a very practical approach (see for example ref. Pich et al., 2019). Full-size genomic EBV DNA, which is approximately 180 kbps in size can be easily purified from E. coli cells but the critical step is reconstitution of infectious virus in a suitable eukaryotic host.

For this, we tested many different eukaryotic cell lines but the only cell that readily gave rise to infectious EBV was HEK293 (human embryonic kidney 293) (Graham et al., 1977). With exceptions, HEK293 cells are still the source of recombinant EBV stocks today (Feederle et al., 2010). After DNA transfection of EBV’s BACmid DNA, cells are selected for resistance against hygromycin or puromycin encoded by *hgy or pac*, respectively, which are cloned onto the prokaryotic mini-F-factor backbone of the BACmid. Selected clonal HEK293 cells establish a latent infection reminiscent of latently EBV-infected cells. To induce virus production, the selected HEK293 clones are transiently transfected with an expression plasmid encoding BZLF1 (Hammerschmidt and Sugden, 1989) to start EBV’s lytic phase (Delecluse et al., 1998; Feederle et al., 2000). This approach is feasible and has become a common quasi-standard among variations on the theme (Feederle et al., 2010; Pich et al., 2019).

Despite this proven approach and reliable source of infectious EBV stocks, it has always been questionable, whether the HEK293 cell is the ideal host to propagate the virus. The nature of this cell line is likely of neuronal origin (Shaw et al., 2002), a cell type never linked to EBV infection or synthesis. Therefore, certain viral genes might be insufficiently expressed such that virus production in HEK293 cells is suboptimal at best. Conversely, certain non-identified viral genes could exist that act as regulators of cellular or viral gene expression dampening or restricting virus production. Such a viral regulator could decrease virus production in cells when expressed in an untimely fashion and/or at inadequately high levels. The identification of such a hypothetical viral regulatory function would also help to design techniques to improve virus yield from HEK293 cells.

As the search for another established cell that supports EBV production has not been successful, we asked whether ectopic expression of individual viral genes during virus synthesis might yield higher yields. Such a case would indicate that this gene is insufficiently expressed during EBV synthesis in the original, HEK293-based producer cells for the B95.8 lab strain of EBV called 2089 (Delecluse et al., 1998). At least one prominent viral example gene is known. The function of a viral glycoprotein encoded by the EBV open reading frame BALF4 was investigated by us back in 2002. Neuhierl et al. found that the composition of infectious EBV was substantially improved when BALF4 was complemented in 2089 EBV producer cells (Neuhierl et al., 2002) indicating that BALF4 encodes a gene product that is limiting during EBV production in HEK293 cells.

In this work we used a basic experimental design with BALF4 as a proven positive control that allows identifying such viral functions. Practically, we generated EB virus stocks from the 2089 EBV producer cell line by ectopic expression of single EBV genes co-transfected with BZLF1 to induce the lytic cycle in these cells. Alternatively, we developed an shRNA technology to repress selected viral transcripts to knockdown certain viral genes of interest during EBV’s lytic phase. The resulting virus stocks were characterized (**Figure 1**) regarding (i) their physical particle concentration using a nanoparticle tracking analysis (NTA) instrument. Bioparticle concentrations were measured in an (ii) Elijah cell-based assay quantitating bound virion particles on the cell surface. Human primary B cells were incubated with engineered EB virions and their (iii) fusogenic characteristics were directly analyzed in a newly developed functional assay. Lastly, we determine the (iv) virus titer of EB virus stocks using Raji cells as target B cells.

**Figure 1.**
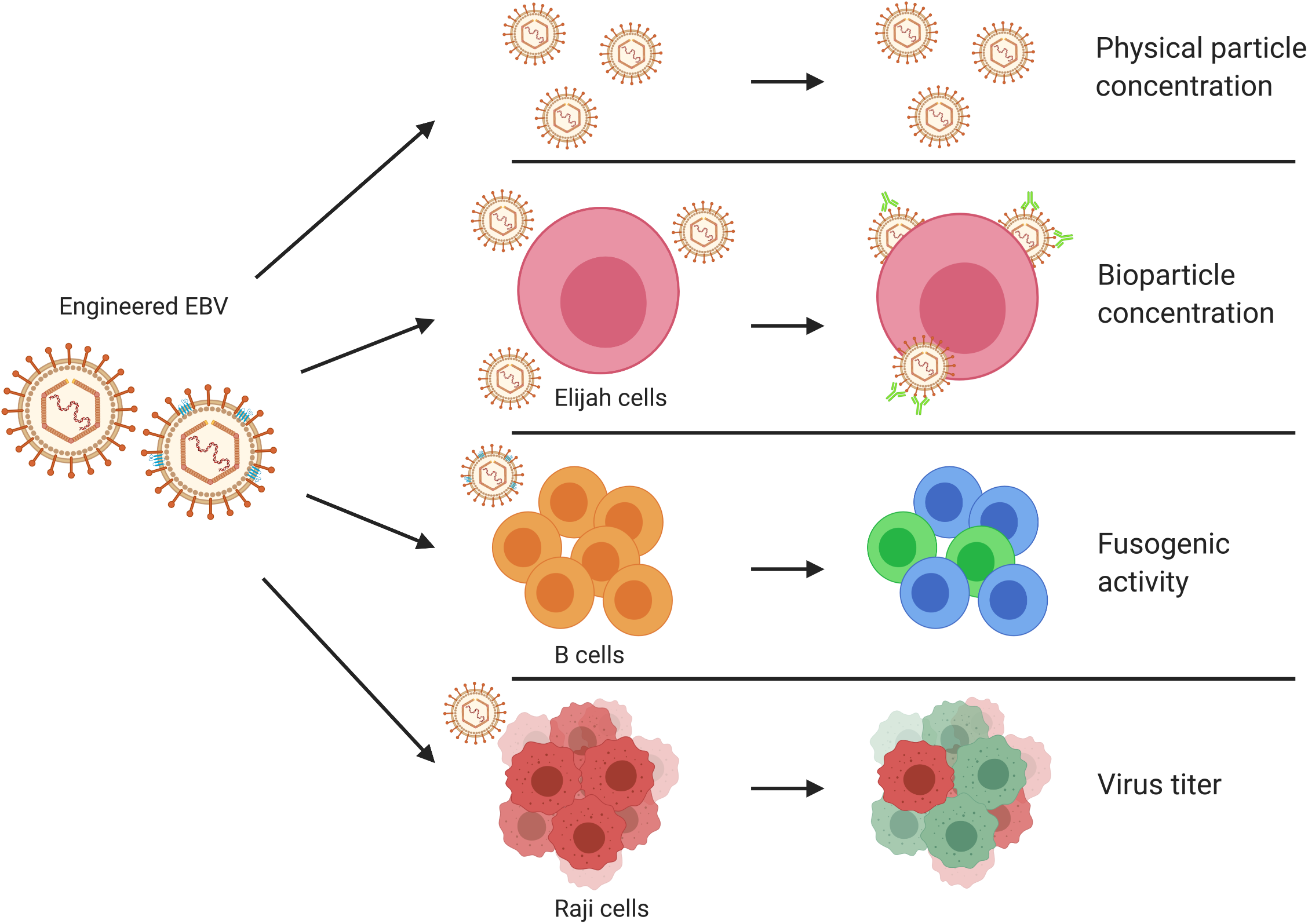
Experimental strategy of analyzing the composition of engineered EBV stocks and their virus-cell interactions. EBV stocks from the EBV producer cell line 2089 were collected three days after transient co-transfection of a BZLF1 expression plasmid together with individual expression plasmids from a panel of 77 different EBV genes. Alternatively, the EBV producer cells were co-transfected with the two plasmids plus a CD63:*ß*-lactamase reporter plasmid to equip virus particles with an enzyme to study viral fusion with target cells. Different individual virus stocks were analyzed for their physical particle concentration using nanoparticle tracking analysis (NTA) instrument, ZetaView PMX110. Elijah cells were used to assess bioparticle concentration in the virus stocks via cell surface binding and subsequent analysis of bound particles by flow cytometry. Virus particles engineered to contain the CD63:*ß*-lactamase reporter protein were analyzed for their fusogenic activity with human primary B cells as targets by flow cytometry. Raji cells were infected with virus stocks to measure the concentration of infectious virus particles conferring green fluorescence protein gene (GFP) in the cells analyzed by flow cytometry. The viral gene designated sLF3 represents a version of LF3 with a reduced number of internal repeats. The diagram was created with the help of BioRender.com (https://biorender.com/).

This comprehensive study surveys 77 viral genes and provides information regarding relevant physical and biological parameter of EBV stocks. The study identifies distinct classes of EBV genes with essential, contributing or dispensable functions regarding these parameters and hints at many viral genes which act dose-dependently. The survey not only has the potential to improve experiments that make use of the maxi-EBV technology but is a rich source of information on lytic phase EBV genes.

## Results

### Establishment of an expression plasmid library with 77 individual viral genes and generation of EBV stocks

Individual open reading frames and known genes of EBV were cloned into the expression vector plasmid pcDNA3 to establish a comprehensive EBV library with all known and hypothetical protein-encoding genes (**Supplementary Table S1**). The purpose of this step was to express single viral proteins ectopically in 2089 EBV producer cells concomitant with inducing EBV’s lytic phase as described (Pich et al., 2019). The plasmid library represents unmodified authentic viral genes without epitope tags that might compromise protein functions. Since there are no specific antibodies available against the majority of EBV’s lytic proteins, the same but epitope-tagged viral genes are contained in a corresponding second plasmid library confirming viral protein expression in HEK293 cells. This way, we can be confident that all unmodified 77 viral genes are also expressed upon their individual transient transfection in 2089 EBV producer cells in this study.

The standard protocol to produce virus stocks is based on the 2089 EBV producer cell line, which are HEK293 cells with a recombinant EBV B95.8 strain genome encoding a green fluorescence protein (Delecluse et al., 1998). Virus production relies on transient transfection of the BZLF1 expression plasmid p509 (Hammerschmidt and Sugden, 1988). To test individual members of the viral expression plasmid library, EBV producer cells were seeded in 6-well cluster plates. After overnight cultivation, the medium was exchanged with 2 ml fresh exosome-depleted (Ex^−^) medium (**Fig. 2**) and the cells were transiently co-transfected with an expression plasmid encoding the BZLF1 gene (p509) together with a plasmid encoding one of the 77 individual viral genes. An ‘empty’ expression vector plasmid was co-transfected with BZLF1 as control as well as reference for subsequent normalization of data. Virus stocks co-transfected with BZLF1 and BALF4 were used as positive controls (Neuhierl et al., 2002). After three days, the virus supernatants were harvested and the resulting virus stocks were tested as described below. All experiments were done in triplicates.

**Figure 2.**
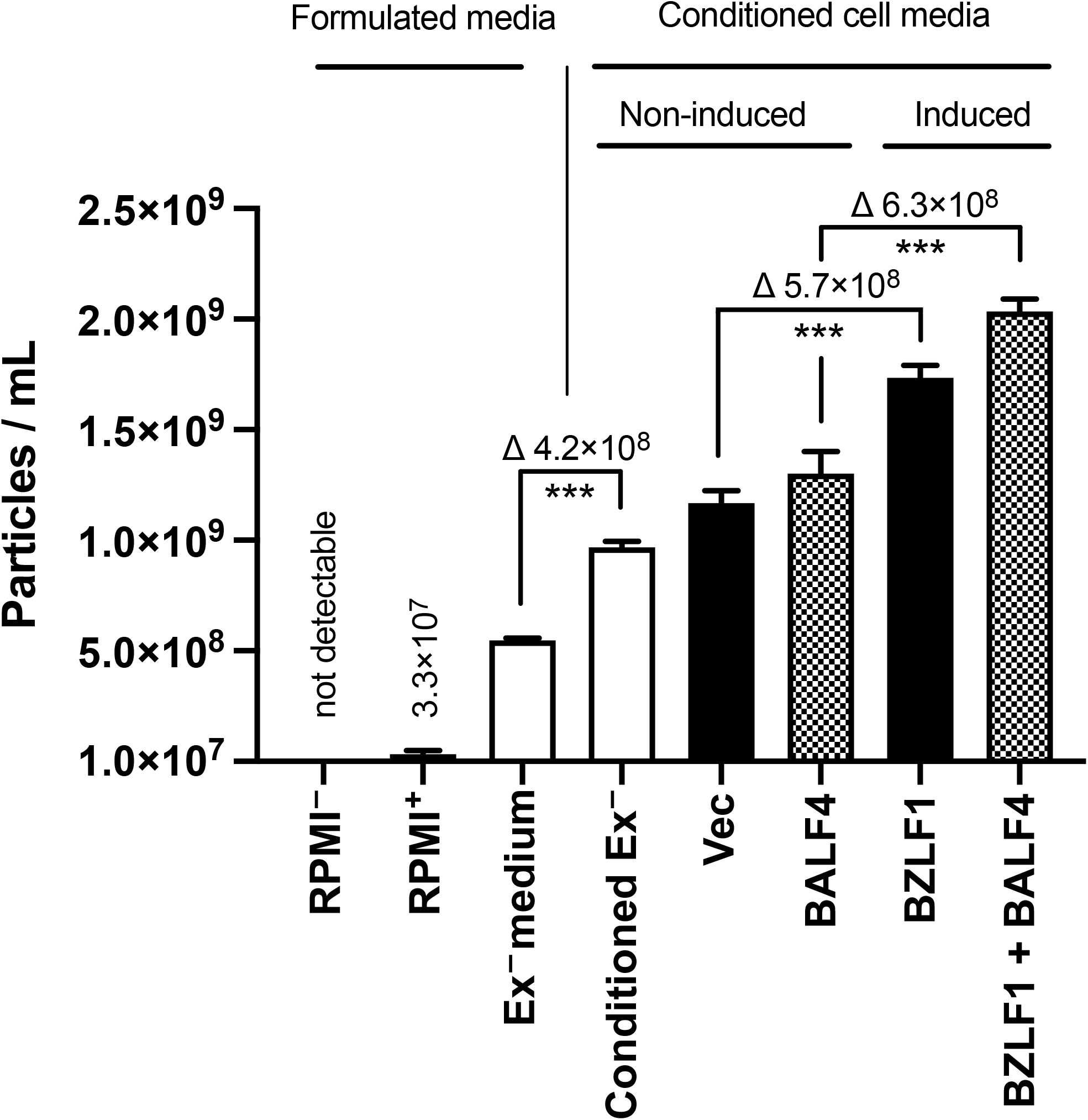
Concentration of physical particles in different formulated cell culture media and in conditioned cell culture media from non-induced and induced 2089 EBV producer cells. Concentrations of physical particles were determined with the aid of the nanoparticle tracking analysis (NTA) using the ZetaView PMX110 instrument. Formulated media are different cell culture medium preparations based on standard RPMI1640, which are as follows: RPMI^−^: plain, non-supplemented commercial RMPI 1640 medium (Gibco); RPMI^+^: commercial RPMI1640 medium supplemented with penicillin-streptomycin stock, sodium pyruvate, sodium selenite, and α-thioglycerols; Ex^−^ medium: RPMI^+^ with 10 % fetal bovine serum and the supplements listed above after ultracentrifugation (100,000 g, 4 °C for more than 16 hours as described in Materials and Methods); Conditioned Ex^−^: Supernatant from non-induced 2089 EBV producer cell line cultured in Ex^−^ medium for three days; Vec: 2089 EBV producer cells transfected with pCMV control plasmid DNA (0.5 μg; p6815) using 3 μl PEI MAX cultivated in Ex^−^ medium for three days; BALF4: same as ‘Vec’ but the cells were transfected with 0.5 μg of the BALF4 expression plasmid p6515; BZLF1: same as ‘Vec’, but the cells were transfected with 0.5 μg of the BZLF1 expression plasmid p509 to induce EBV production; BZLF1 + BALF4: same as ‘Vec’, but the cells were co-transfected with 0.25 μg each of the expression plasmids p509 and p6515 coding for BZLF1 and BALF4, respectively. Mean and standard deviation of three replicates are shown. *** P≤0.001.

### Quantitative analysis of extracellular vesicles (EVs) in cell culture media from non-induced and induced 2089 EBV producer cells

Standard cell culture medium contains fetal bovine serum, which introduces abundant quantities of extracellular vesicles and related particles in the range of 50 to more than 200 nm. We analyzed the concentration of EVs in plain RPMI1640, formulated cell culture medium based on RPMI1640 as well as in conditioned cell culture supernatants using a ZetaView PMX110 instrument capable of nanoparticle tracking analysis (NTA). The results are shown in **Figure 2**. Fully supplemented but partially EV-depleted RPMI1640 cell culture medium contains approximately 5×10^8^ physical particles per ml, which are of bovine origin (**Fig. 2**). This formulated cell culture medium was termed ‘Ex^−^ medium’ in which non-induced 2089 EBV producer cells were cultivated for three days. This ‘Conditioned Ex^−^ medium’ contained considerably more EVs (**Fig. 2**). Transient transfection of two plasmid control DNAs (indicated ‘Vec’ and ‘BALF4’ in **Fig. 2**) followed by cultivating the non-induced cells for three days increased the EV concentration further. Finally, EBV’s lytic phase was induced by transfecting expression plasmids encoding BZLF1 only or in combination with BALF4, which caused a significant increase of physical particles of 5.7×10^8^ and 6.3×10^8^ EVs per ml, respectively, compared to the corresponding non-induced controls after three days (**Fig. 2**).

This analysis documents that conventional cell culture medium contains substantial numbers of EV even after depleting bovine EVs. Spent, conditioned medium contains more EVs, which all cells spontaneously release. After lytic induction of 2089 EBV producer cells, EV concentrations increased further suggesting that induced, virus-releasing cells shed more physical particles (in the order of 5×10^8^ per ml) than non-induced cells.

### Effects of individual EBV genes on physical particle concentration

To test all 77 individual members of the viral expression plasmid library, 2089 EBV producer cells (Delecluse et al., 1998) were transiently co-transfected with plasmid DNAs encoding the BZLF1 gene and one of the 77 individual viral genes. BALF4 was used as a positive control. Three days after transfection the virus supernatants were investigated for their concentrations of physical particles by NTA. **Figure 3** shows the results of 77 viral genes plus controls in descending order. Mean particle concentration in the different virus stocks varied in between factors of 0.7 to 1.3. Certain virus stocks showed inconsistent numbers of physical particles from one experiment to the next documented by their wider standard deviation (e.g., BcLF1, BVRF1, BGLF3.5, LF1 among others) for unknown reasons.

**Figure 3.**
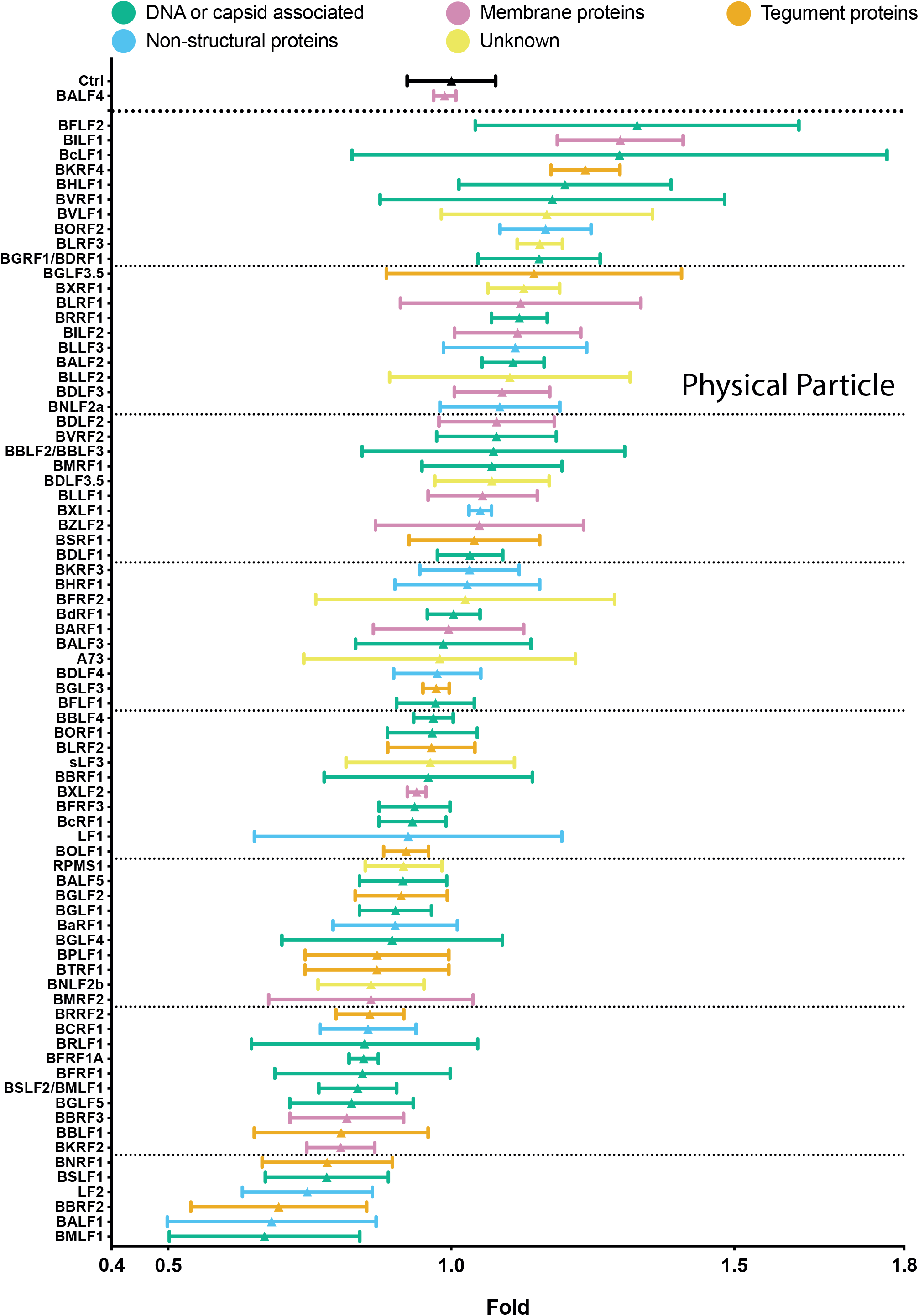
Comparison of physical particle concentrations in virus samples generated by co-transfection of BZLF1 and single expression plasmids from a panel of 77 EBV genes including two controls. EB virus stocks were analyzed for their physical particle concentration by nanoparticle tracking analysis (NTA). NTA was performed with the ZetaView PMX110 instrument and the images were analyzed with the ZetaView 8.04.02 software. Standard calibration beads were used to confirm the range of linearity. The y-axis lists the BZLF1 co-transfected individual EBV genes. An empty pCMV vector plasmid plus the BZLF1 plasmid were co-transfected as reference (Ctrl). The positive control encompasses supernatants obtained after co-transfection of both BZLF1 and BALF4 expression plasmids. The number of physical particles in the range of 100 – 200 nm contained in the supernatants of cells were analyzed and normalized to the reference sample (Ctrl). The results are arranged in descending order and are classified according to five functional groups and color-coded as indicated. Mean and standard deviation of three biological replicates are shown. The horizontal lines indicate groups of 10 viral genes for better visualization. The viral gene designated sLF3 represents a version of LF3 with a reduced number of internal repeats.

We hypothesized that there is a link between physical particle numbers and molecular functions of the genes tested. Similarly, we wondered if their expression timing could correlate with the results. We also wanted to select interesting genes to proceed. For this purpose, we depicted the data in a color-coded fashion in **Figure 3** and **Supplementary Figure S1**. We selected ten genes at the high end, five of which are associated with DNA or capsid functions, one encodes a membrane protein, a tegument protein, a non-structural protein and two have unknown functions. At the low end are 16 genes, seven of which are associated with DNA or capsid functions, two encode membrane proteins, four encode tegument proteins and three are non-structural proteins. Regarding expression timing, within the top 10 genes, 2 are expressed early, 4 are expressed late and the remaining have no known expression timing (**Supplementary Fig. S1**). In the group of 16 virus stocks, which have the lowest number of physical particles, 7 are expressed early, 5 are expressed late and the expression timing of the remining genes is unknown. No particular function appears to cluster in the virus stocks arranged in descending order. Also, expression timing does not reveal an obvious pattern suggesting that various viral genes expressed throughout the entire lytic phase of virus production and egress can modulate the release of physical particles as measured by the NTA instrument.

### Measuring viral bioparticle concentrations

Viral infectivity is the successful endpoint of the infectious route, which initiates with virus binding to targets cells as the very first step. We measured this key function using a cellular binding assays with Elijah cells as target, a human B cell line derived from a case of Burkitt’s lymphoma (Rowe et al., 1985). The binding assay, which to our knowledge was introduced in 2010 (Busse et al., 2010) is a proxy of virus attachment to human B cells, which includes EBV’s cognate CD21 receptor, the complement receptor 2 (CR2) (Nemerow et al., 1987; Tanner et al., 1987) and the viral gp350 protein encoded by BLLF1. It has remained uncertain, whether binding of EBV particles to the surface of Elijah cells solely depends on the interaction of gp350 with CD21 or whether additional viral glycoproteins contribute to surface binding as suggested earlier (Janz et al., 2000). Therefore, we deleted *CD21* using a CRISPR-Cas9 technology (Akidil et al., 2021) in Elijah cells and found that EBV binding was no longer detectable even with exceedingly high virus doses (**Fig. 4B**). This figure shows that binding of viral particles to Elijah cells is solely dependent on CD21, the initial receptor of EBV required for viral adhesion to B cells (Fingeroth et al., 1984; Nemerow et al., 1985, 1987, 1989). Our finding also indicates that the Elijah cell binding assay exclusively records gp350 and gp350-bearing particles, which include intact virions but also non-infectious particles such as defective viruses, virus-like particles and gp350 decorated extracellular vesicles (EVs), which are released from lytically induced 2089 EBV producer cells. As the Elijah cell binding assay cannot differentiate between various gp350 containing vesicle species we collectively call them ‘bioparticles’ throughout this study.

**Figure 4.**
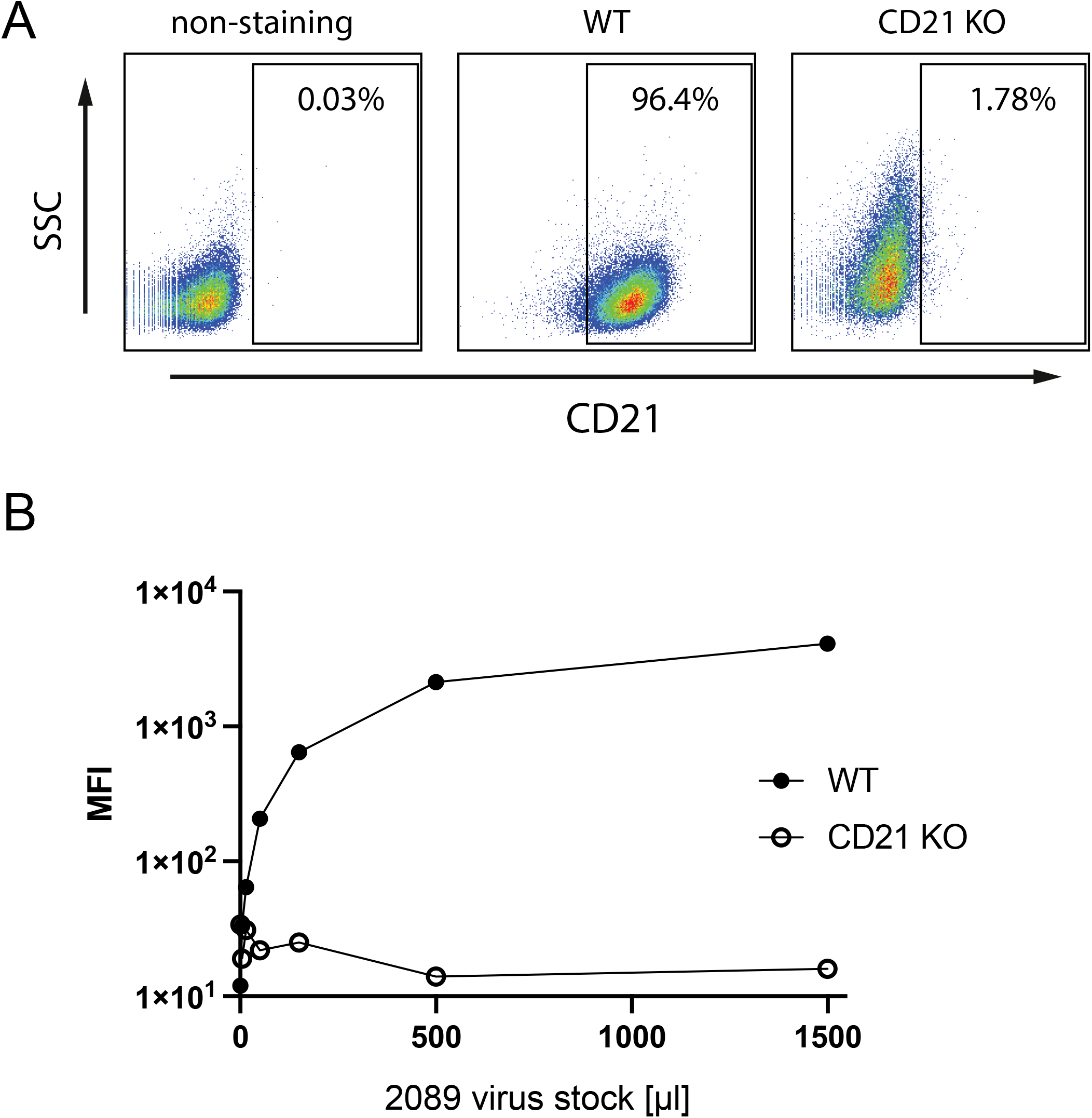
CD21 is essential for virus-cell adhesion. The two alleles of the surface receptor CD21 were deleted in Elijah cell chromatin using preformed CRISPR-Cas9 ribonucleoprotein (RNP) complexes with two individual CD21-targeting guide RNAs (gRNA) and a recombinant Cas9 nuclease. The cells were further sorted for CD21 negative cells. **(A)** CD21 expression levels and knock-out efficiency of Elijah cells were evaluated by flow cytometry using an APC-coupled CD21-specific antibody. Left panel: wild-type Elijah cells without antibody staining; middle: wild-type Elijah cells (WT) stained with the fluorochrome coupled CD21 antibody; right: CD21 expression level in sorted CD21 negative Elijah cells after CRISPR-Cas9 mediated knockout (CD21 KO). **(B)** Binding activity of 2089 EB virus stocks was analyzed with wild-type (WT) and CD21 knock-out (CD21 KO) Elijah cells. 2×10^5^ Elijah cells were incubated with 0, 5, 15, 50, 150, 500 and 1500 μl virus stocks at 4 °C for 3 h. The cell-surface-bound virus particles were detected with an Alxea647-coupled anti-gp350 antibody, mean fluorescence intensities (MFI) were recorded by flow cytometry and plotted as a function of virus dose.

The previously tested EBV virus stocks and appropriate controls shown in **Figure 3** were analyzed for bioparticle binding to the surface of Elijah cells. Practically, Elijah cells and individual virus stocks were incubated for a limited time at low temperature to allow firm binding to Elijah cells. Binding was detected with the gp350-specific, fluorochrome-coupled monoclonal 6G4 antibody. After appropriate washing steps the cells were analyzed by flow cytometry for their raw values of mean fluorescence intensity (MFI). Virus stocks generated with the 2089 EBV producer cells after co-transfection of an empty vector plasmid together with BZLF1 (p509) served as the reference control (Ctrl) and standard for calculating the ratios of MFI values as shown in **Figure 5** and **Supplementary Figure S2**. Virus stocks generated by co-transfection of the BALF4 encoding expression plasmid p6515 together with p509 served as an additional control as the number of viral particles obtained after ectopic expression of BALF4 was expected to be unaltered (Neuhierl et al., 2002).

**Figure 5.**
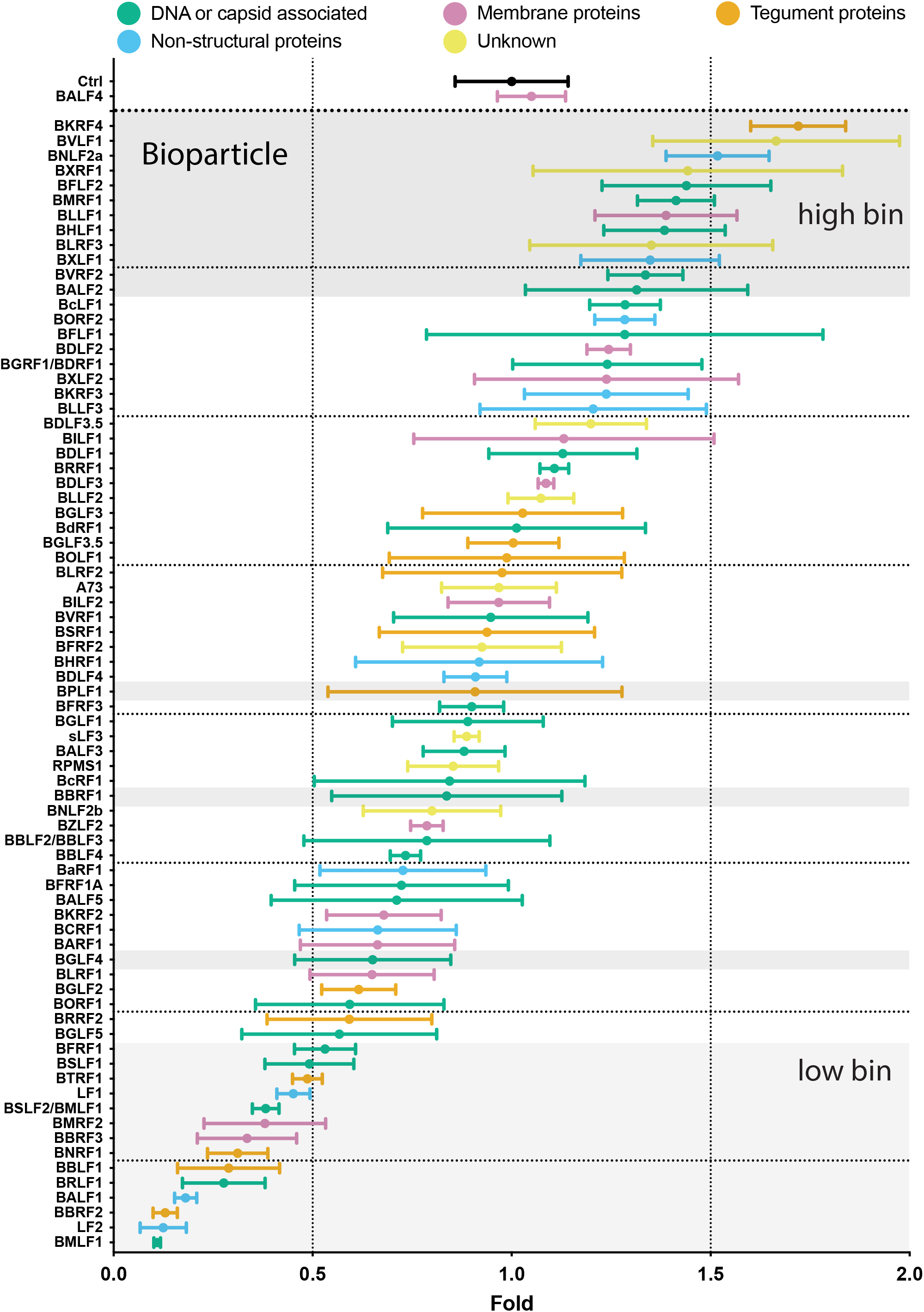
Comparison of bioparticle concentrations in virus samples generated by co-transfection of 2089 EBV producer cells with BZLF1 and individual expression plasmids from a panel of 77 EBV genes. Virus production and generation of samples were identical as described in the legend of **Figure 3**. The amount of gp350 positive particles bound to Elijah cells was quantified using a gp350-specific, fluorochrome-coupled monoclonal antibody and flow cytometry. The ratios of mean fluorescence intensity (MFI) values of individual samples versus the MFI value of the reference sample (Ctrl) were calculated and are provided on the x-axis. An empty pCMV vector plasmid co-transfected with the BZLF1 plasmid p509 served as reference (Ctrl). The y-axis lists the transfected individual EBV genes. Ratios are arranged in descending order. An expression plasmid encoding BALF4 served as a positive control. Shaded areas at the top and bottom highlight groups of viral genes termed ‘high bin’ and ‘low bin’, respectively. Three singly highlighted genes (BPLF1, BBRF1, BGLF4) were randomly picked as further candidates. EBV genes are classified according to five functional groups as indicated. Mean and standard deviation of three biological replicates are shown. The two vertical lines indicate 0.5- and 1.5-fold ratios, horizontal lines indicate groups of 10 viral genes for better visualization. The viral gene designated sLF3 represents a version of LF3 with a reduced number of internal repeats.

In fact, virus stocks produced by co-transfecting BALF4 and BZLF1 showed absolutely no difference in this assay (**Fig. 5**). Other virus stocks showed a similar range of variability of about a factor of 1.5 to 0.5 as already seen in **Figure 3**. Only a small group of 13 virus stocks (BSLF1 -> BMLF1) showed a reduction below a factor of 0.5. Identical to **Figure 3**, the color codes illustrate the five functional groups of viral proteins in **Figure 5**. Again, no discrete functional characteristics was obvious. When the results were color-coded according to gene expression timing regarding early, late genes or genes with unknown timing, no special contribution of the three different classes could be found (**Supplementary Fig. S2**). This result suggests that various viral genes expressed throughout the entire lytic phase of virus production contribute to viral bioparticles synthesis but clearly certain viral genes can increase, reduce or even repress it.

BKRF4, BVLF1 and BNLF2a, which represent a tegument protein, a protein with unknown function and a non-structural protein, respectively, are top candidates, which together with seven additional candidates from a group which we designated ‘high bin’ (**Fig. 5**). BKRF4, BVLF1 and BNLF2a all belong to different groups regarding expression timing (**Supplementary Fig. S2**), but the three genes showed a similar effect and increase bioparticle production by about 50 % compared to the control. Fourteen viral genes that decreased bioparticle concentration by a factor of approximately 0.5 (BFRF1, BSLF1, BTRF1, LF1, BSLF2/BMLF1, BMRF2, BBRF3, BNRF1, BBLF1, BRLF1, BALF1, BBRF2, LF2, BMLF1) and were designated members of the ‘low bin’ group (**Fig. 5**). Five genes among this class belong to DNA or capsid-associated proteins, two are membrane proteins, four are tegument proteins and three belong to the group of non-structural proteins. Based on expression timing, six genes are expressed early, five are expressed late and three have no known expression timing. Remarkably, virus stocks generated by ectopic expression of BALF1, BBRF2, LF2 and BMLF1 genes have a considerably reduced viral bioparticle concentrations in the order of 0.1, only (**Fig. 5**).

Similar to effects of individual EBV genes on physical particle concentration, they also modulate the release of bioparticles, but a coherent picture is not apparent.

### Ectopic expression of certain viral genes increases or decreases virus titers, but BALF4 is unique

To analyze the infectivity of the virus stocks, we used Raji cells, a human Burkitt’s lymphoma cell line (Pulvertaft, 1964), which is a proven model. The fraction of EBV-infected Raji cells was identified by green fluorescence protein expression and quantified by flow cytometry. The results from three independent infection experiments were normalized to the respective controls, which were set to one. The ratios of the 77 tested combinations plus controls were arranged in descending order and color-coded as before (**Fig. 6** and **Supplementary Fig. S3**). The viral BALF4 gene co-transfected with BZLF1 served as a positive control. The results in **Figure 6** revealed that ectopic expression of BMRF1, BGRF1/BDRF1, LF1, BLRF3, BDLF2 or BDLF3.5 yielded virus titers increased by a factor of 1.5 or more indicating that only six genes besides BALF4 support higher virus titers. Two genes encode DNA or capsid-associated proteins, one is a membrane protein, one is a non-structural protein, and two genes have unknown functions. No tegument protein encoding gene is among this list. Expression timing was not informative, either (**Supplementary Fig. S3**). The majority of viral genes tested, about 60 in total, are in a neutral zone with little variations to virus titers at around control level. In contrast, the last 15 expression plasmids (BGLF5, BTRF1, BBRF1, BFRF1, BSLF2/BMLF1, BBRF3, BNRF1, BBLF1, BRLF1, BMRF2, BGLF4, BBRF2, BMLF1, LF2, BALF1) co-transfected with BZLF1 led to virus titers reduced by a factor of 2 and more (**Fig. 6**). Again, there is no clear attributable functionality nor does expression timing seems to be a notable criterion.

**Figure 6.**
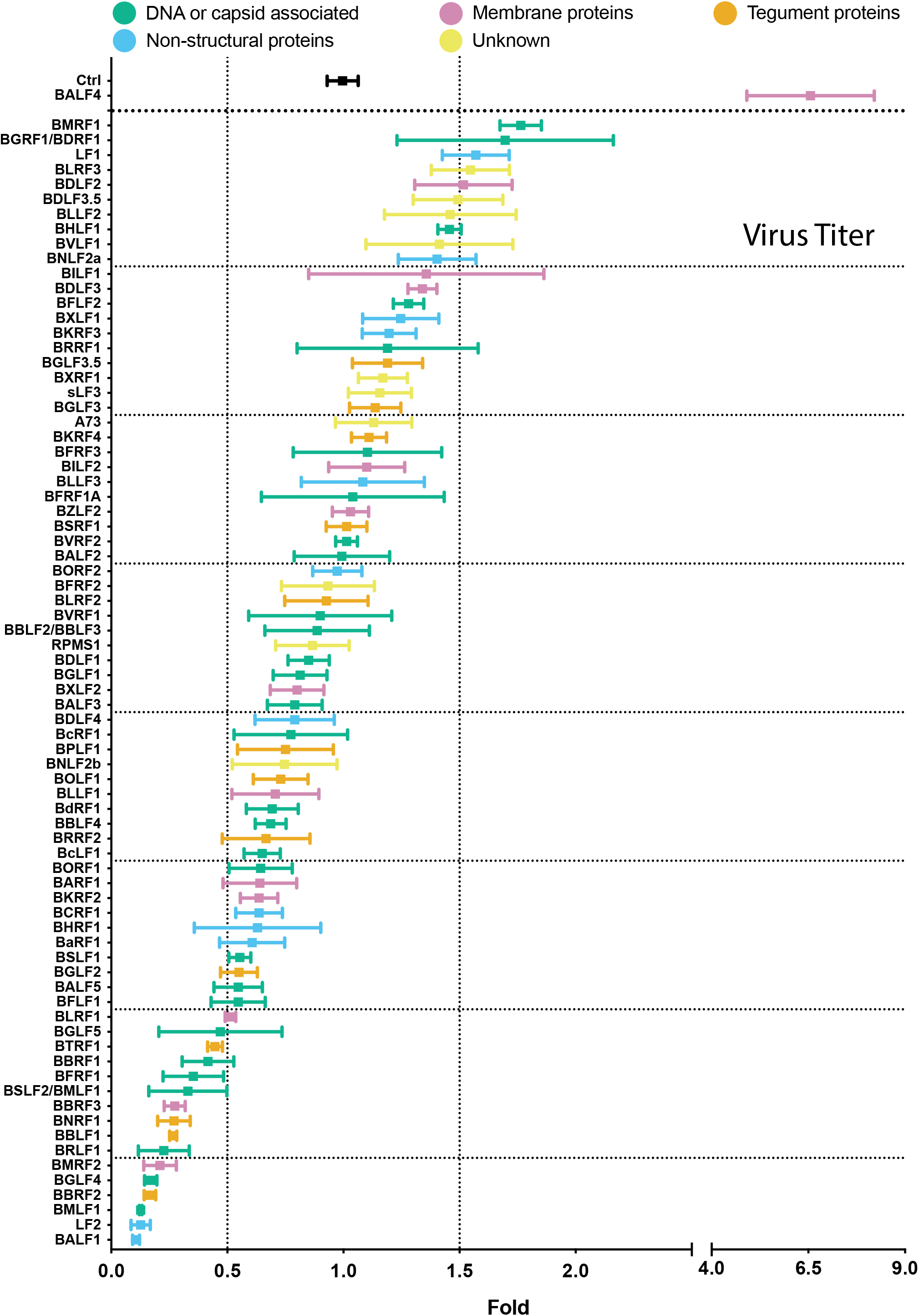
Comparison of virus titer of stocks generated by co-transfecting 2089 EBV producer cells with BZLF1 together with single expression plasmids from a panel of 77 EBV genes. 2089 EBV producer cells were co-transfected with BZLF1 and single expression plasmids encoding the denoted viral genes. Virus containing supernatants were harvested 3 days later and used to infect Raji cells. After three days, the infected Raji cells were investigated by flow cytometry analyzing the expression of green fluorescence protein. The y-axis lists the individual EBV genes transfected in combination with BZLF1. The BZLF1 (p509) expression plasmid co-transfected with p6816, an empty pCMV vector plasmid served as reference and control (Ctrl). The titers of infectious EBV stocks according to the percentage of GFP-positive Raji cells (GRU) were normalized to the reference sample (Ctrl), which was set to 1. The results are listed in descending order. EBV genes are color-coded according to five functional groups as indicated. Mean and standard deviation of three biological replicates are shown. Vertical lines indicate 0.5- and 1.5-fold ratios. The horizontal lines indicate groups of 10 viral genes for better visualization. The viral gene designated sLF3 represents a version of LF3 with a reduced number of internal repeats.

Taken together, ectopic expression of individual viral genes resulted in small differences in the concentration of the virus stocks, only. The positive control BALF4 is clearly superior to the remaining viral genes tested and increases the infectivity of the virus stocks by a factor of 6.5, which exceeds the effects of other viral genes by far (**Fig. 6**). Remarkably, the B95.8 laboratory strain lacks the viral LF1 gene suggesting that it contributes a previously unknown supportive functions during virus synthesis or enhances viral infectivity. A larger group of genes exhibits repressive functions and shows a clear negative impact on virus synthesis or virus infectivity.

### Statistical analysis reveals a solid correlation of physical particle and bioparticle concentrations and virus titers

The findings shown in **Figures 3** to **6** document the characteristics of three quantitative parameters (physical particle and bioparticle concentrations, virus titers) obtained from 77 individual EBV stocks. The mean values of the three measurements of the virus stocks were taken and set as rank order to investigate a possible correlation using the Spearman rank correlation method. The values are provided in a heatmap shown in **Supplementary Figure S4** and show a high overall correlation.

To look for correlative details in the data, mean values derived from the three quantitative parameters were sorted according to viral functions or grouped according to expression timing criteria. No statistically significant contribution of viral proteins with different functions or different expression timing was found to contribute to the three quantitative parameters analyzed (**Supplementary Figure S4**).

The statistical analysis suggests that individual viral genes can have a solid impact on all three basic parameters investigated. No correlations regarding expression timing or functional classes emerge demonstrating that HEK293 cells seem to support virus synthesis in general with no ‘sweet spot’ identified so far except BALF4.

### Selection, design and validation of shRNA candidates targeting 25 viral genes

To concentrate on potentially interesting viral genes, we chose candidates from both ends of the ranked candidates identified in the experiments shown in **Figure 5** and asked if members of the ‘high bin’ group are essential for virus production as measured in infection experiments in **Figure 6**. Conversely, we asked if members of the ‘low bin’ group of viral genes might regulate virus synthesis negatively and hence repress virus titers, because certain genes might act as master regulators controlling and thus limiting virus production in HEK293 cells.

As shown in **Figure 5**, the ‘high bin’ group encompasses ten candidates (BKRF4, BVLF1, BNLF2a, BXRF1, BFLF2, BMRF1, BHLF1, BXLF1, BVRF2, BALF2) and the ‘low bin’ group lists eleven viral genes (BFRF1, BSLF1, BTRF1, BMRF2, BBRF3, BNRF1, BBLF1, BRLF1, BALF1, BBRF2, BMLF1). Three randomly selected genes, BPLF1, BBRF1 and BGLF4, were chosen from the large middle group. Certain genes were excluded from the ‘top bin’ and ‘low bin’ groups for various reasons. LF1 and LF2 were excluded since the B95.8 EBV strain in the 2089 EBV producer cell line does not encompass these two genes. Their ectopic expression in the 2089 producer cells was informative but their lack precludes a knockdown strategy as explained below. BLLF1 (gp350) was excluded because its ectopic expression directly affected the quantitative analysis of bioparticles. Also, BLRF3 was excluded from the ‘high bin’ group as it is part of EBNA3 and likely not directly involved in EBV’s lytic phase.

To test the 25 selected viral genes for their possible essential or regulatory roles during lytic infection, we used an shRNA knockdown strategy to repress individual viral gene products to study their contribution during virus and bioparticle production. The design of the shRNAs and their functional evaluation are shown in **Supplementary Figure S5** and are described in the Materials and Methods section.

### Virus titers in supernatants from 2089 EBV producer cell lines stably transduced with sets of shRNAs directed against selected EBV transcripts

Twenty-four EBV producer cell lines stably transduced with sets of three shRNA vectors each were established under co-selection with puromycin and hygromycin B to ensure shRNA expression and maintenance of the EBV 2089 genome, respectively. Controls included 2089 EBV producer cells transduced with the empty shRNA vector backbone (p6924), only, as well as cells encoding triple sets of shRNAs directed against GFP or BALF4 transcripts as tested in **Supplementary Figure S5**. The three graphs in **Figure 7A** depict the selected genes, their functional attributes and recapitulate bioparticle ratios obtained after the genes’ ectopic expression shown in **Figure 5**. Virus stocks from the different 2089 EBV producer cell lines stably transduced with shRNA vectors were generated by transient BZLF1 transfection and quantified by infecting Raji cells as in **Figure 6**. The results were arranged according to the ‘high bin’ and ‘low bin’ groups in **Figure 7B** (red columns) and shown in comparison with ratios of virus titers when the viral genes were ectopically expressed in the parental 2089 EBV producer cell line (black columns).

**Figure 7.**
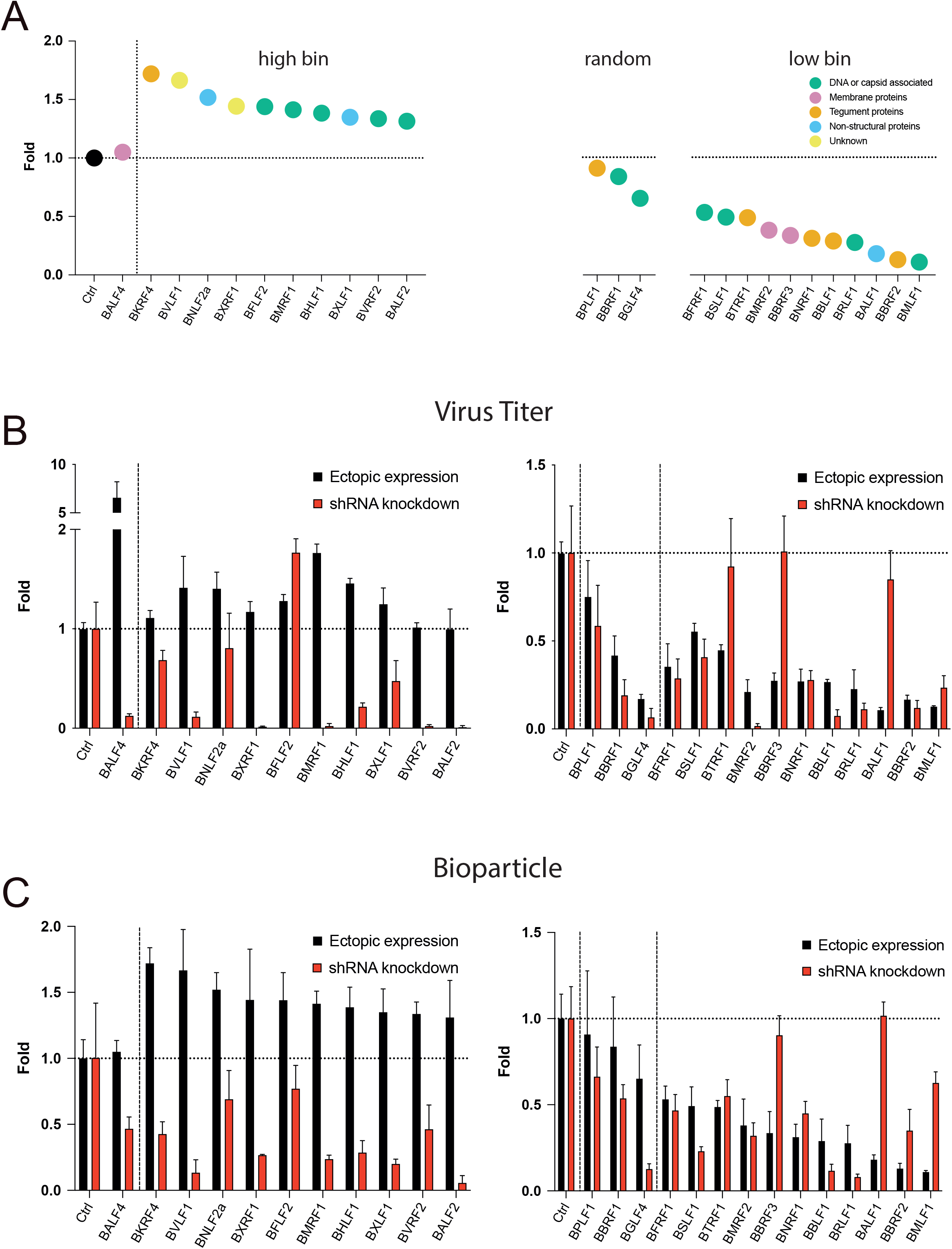
Viral titers and bioparticles in supernatants generated after co-transfection of BZLF1 and single viral genes into 2089 EBV producer cell lines stably transduced with sets of three shRNAs directed against 25 individual viral transcripts. **(A)** Recapitulation of selected data shown in **Figure 5**. Three groups of viral genes are shown that were picked randomly (’random’), or increased (’high bin’, left panel) or decreased (’low bin’, right panel) the yield of bioparticles when co-transfected together with BZLF1 into the EBV producer cell line 2089. The EBV genes are arranged in descending order and the color codes depict the different functions of viral proteins as in Figure 5. **(B)** Parental 2089 EBV producer cells were stably transduced with sets of three shRNAs each that target 25 viral transcripts. Controls encompass the EBV producer cell line 2089 stably transduced with the empty pCDH shRNA expression vector (Ctrl). The viral genes termed ‘high bin’ and ‘low bin’ groups are shown in the left and right panels, respectively. The individual cell lines were transiently transfected with the BZLF1 expression plasmid. The virus supernatants were harvested 3 days after transfection and used to infect Raji cells. After three days the fraction of GFP-positive Raji cells was determined by flow cytometry and virus titers (GRU titers) were calculated. Supernatants harvested from the control cell lines are shown on the left side of the graph with the ‘high bin’ group of genes separated by a vertical dotted line from the test samples. The ‘low bin’ group is shown on the right separated from the control and includes the three ‘random’ genes separated by a dotted vertical line. Virus titers of supernatants obtained from the 25 individual shRNA expressing EBV producer cell lines (red columns; shRNA knockdown) analyzed in panel B are compared with virus titers found in supernatants from the parental EBV producer cell line 2089 upon transient expression of viral genes (black columns; ectopic expression). Mean and standard deviation of three biological replicates are shown. **(C)** Virus stocks analyzed in panel B were tested for their bioparticle concentration in the Elijah cell binding assay. Bioparticle concentrations found in supernatants from parental 2089 EBV producer cells upon ectopic expression of single viral genes (as analyzed in Figure 5; black columns; ectopic expression) are compared with results obtained from 2089 EBV producer cells stably transduced with sets of shRNAs (red columns; shRNA knockdown). Mean and standard deviation of three biological replicates are shown.

In the left panel of **Figure 7B**, ectopic expression of BALF4 yielded superior virus titers whereas its knockdown repressed virus titers to about 10 %. These results validated the shRNA knockdown strategy and confirmed BALF4’s role as an essential gene (Neuhierl et al., 2009) that boosts viral infectivity upon ectopic expression (Neuhierl et al., 2002).

As expected, certain shRNA sets decreased virus titers dramatically but not all shRNA sets caused a serious phenotype (**Fig. 7B**). The results indicated that certain viral genes seem to be absolutely essential (BXRF1, BMRF1, BVRF2, BALF2, BMRF2) or, conversely, are entirely dispensable (BFLF2, BTRF1, BBRF3 and BALF1) for the production of infectious virus stocks.. Other shRNAs had a weak repressive phenotype (BKRF4, BNLF2a, BHLF1, BXLF1, BFRF1, BSLF1, BBLF1, BRLF1) or hardly affected production of infectious virus (BPLF1). Three shRNAs (BTRF1, BBRF3, BALF1) in the ‘low bin’ group (**Fig. 5**) did not increase virus production beyond wild-type level suggesting that the three genes also do not qualify as regulators of EBV’s lytic phase although their ectopic expression was detrimental for the release of infectious EBV (right panel in **Fig. 7B**). Remarkably, the shRNA mediated knockdown of BFLF2 seemed to induce virus titers to some degree (**Fig. 7B**, left panel).

### Ectopic expression and shRNA knockdown of viral genes and transcripts – comparing virus titers and bioparticle concentration of 25 viral targets

Virus stocks from infection experiments with Raji cells (**Figure 7B**) were also analyzed for their bioparticle concentrations using the Elijah cell binding assay. The results are summarized in **Figure 7C**. Results from ectopic expression of single viral genes and the matching shRNA knockdown experiments are shown in black and red, respectively. The shRNA mediated knockdown of BALF4 showed a decrease by half of bioparticle concentration, but a substantial reduction in the BVLF1 and BALF2 shRNA expressing EBV producer cell lines was observed, which is in agreement with their low virus titers in **Figure 7B**. In the right panel of **Figure 7C**, the majority of supernatants from shRNA expressing EBV producer cell lines did not show very different bioparticle concentrations (BBRF1, BFRF1, BTRF1, BMRF2, BNRF1) whereas some had reduced numbers of bioparticles (BGLF4, BSLF1, BBLF1, BRLF1). On the contrary, the shRNA mediated knockdown of BBRF3 and BALF1 transcripts had no measurable effect (**Fig. 7C**, right panel). The 2089 producer cell line transduced with the set of three shRNA directed against the GFP transcript did not affect bioparticle concentration much validating our general approach.

Together, the shRNA strategy provides critical information complementing results from ectopically expressed single viral genes regarding bioparticle concentrations and infectious virus titers in **Figures 5** and **6**, respectively. However, the shRNA strategy failed to identify a viral master regulator, because a knockdown of viral transcripts in the ‘low bin’ group did not increase bioparticle or virus synthesis beyond control levels.

### Analysis of the fusogenic activity of 25 virus stocks from the ‘high bin’ and ‘low bin’ groups

We developed a novel assay to measure the fusogenic activity of engineered extracellular vesicles (EVs) (Albanese et al., 2020), which we adapted recently to analyze the fusogenicity of EB virions using primary human B cells, EBV’s primary target cells, and plasmacytoid dendritic cells, pDCs (Bouvet et al., 2021). The *β*-lactamase (BlaM) assay has been developed in the HIV field to detect fusion of HIV particles with T cells (Cavrois et al., 2002; Jones and Padilla-Parra, 2016). In this context, the enzyme is introduced into recipient cells by virus-mediated transfer to cut the **ß**-lactam ring of CCF4 substrate with which the cells are loaded. Cleavage of the CCF4 substrate alters its emission wavelength from 520 nm to 447 nm, which can be detected and differentiated by flow cytometry.. The protocol has been constantly improved (Cavrois et al., 2014) and the method is applicable to detect transduction of BlaM when incorporated into viruses or other entities such as EVs (Albanese et al., 2020; Bouvet et al., 2021).

To study virus fusion separately and independent of viral infectivity (which encompasses intracellular transport and *de novo* expression of viral proteins or the GFP reporter), the assay shown schematically in **Figure 8A** was established. 2089 EBV producer cells were co-transfected with an expression plasmid encoding CD63:BlaM (p7200) together with BZLF1 (p509) and single expression plasmids that encode the 21 individual viral genes of the ‘high bin’ and ‘low bin’ groups plus controls as defined in **Figures 5** and **7A**. Human primary B cells isolated from adenoid tissue were incubated with CD63:BlaM-containing virus stocks for 4 h. Subsequently, the cells were loaded with CCF4 substrate and incubated at room temperature for 16 h. The fraction of cells with emission light shift was determined by flow cytometry. Virus stocks generated by co-transfection of BZLF1, an empty expression vector plasmid and CD63:BlaM served to normalize the data depicted in **Figure 8B**. The CD63:BlaM equipped virus stocks were also used to determine their titers after Raji cell infection (**Supplementary Fig. S6**). A virus stock generated by transient transfection of the 2089 EBV producer cells with only BZLF1 (p509) served as additional control.

**Figure 8.**
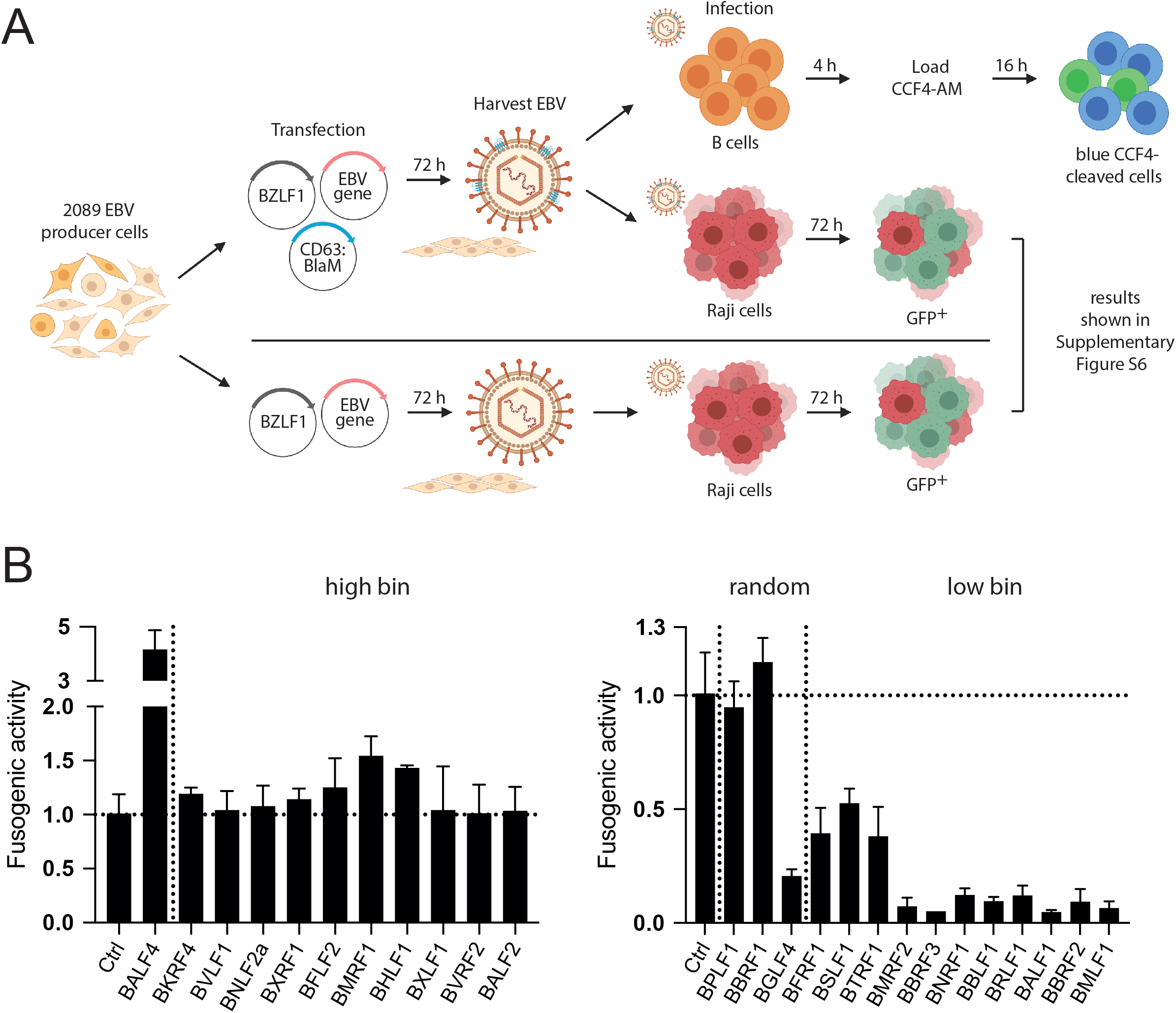
Analysis of engineered virus stocks using the BlaM fusion assay and primary human B cells as targets. **(A)** The flow chart depicts the experimental steps of the fusion assay (top pathway) and its comparison with the Raji cell-based test for infectivity (below, the results are shown in Supplementary Fig. S6). **(B)** Results of the *ß*-lactamase fusion assay with supernatants of the 2089 EBV producer cell line transiently transfected with 25 individual expression plasmids encoding viral genes of the ‘high bin’, ‘low bin’ groups and three random genes. The cells were co-transfected with three plasmids as shown in panel A (plasmids encode BZLF1 [p509], CD63:BlaM [p7200] and one of the 25 selected EBV genes or controls). The resulting supernatants were tested on primary human B cells as targets. The read-outs are based on the fraction of ‘blue’ B cells with cleaved CCF4 substrate by flow cytometry and normalized to the control. Shown are the results from the ‘high bin’ and ‘low bin’ groups and three randomly picked viral genes separated by a vertical dotted line. The illustration was created using the BioRender.com source (https://biorender.com/).

The left panel in **Figure 8B** shows the fusogenic activities of 10 EBV stocks of viral genes (plus BALF4) that belong to the ‘high bin’ group as defined in **Figures 5** and **7A**. The BALF4 virus stock showed a four-fold increase in this assay and another virus stock generated with BMRF1 showed a slightly improved fusogenic activity. Most other virus stocks did not reveal changes in their fusogenic activities. Within the members of the ‘low bin’ group of genes (**Fig. 8B**, right panel), all virus stocks showed medium (BFRF1, BSLF1, BTRF1) or much reduced fusogenic activities (BMRF2 to BMLF1) probably following a similarly reduced bioparticle concentration (**Fig. 5**) in this group of genes. Interestingly, while the fusogenicity of the BGLF4 virus stock was clearly compromised, the two virus stocks belonging to the ‘random’ group, BPLF1 and BBRF1, had comparable fusion activity at the level of the control stock (**Fig. 8B**, right panel). The BBRF1 virus stock is an exception, because its infectivity is reduced by more than 50 % (**Fig. 6**) while its fusogenic activity is wild-type.

Taken together, the assay demonstrates that BALF4 is the dominant driver of viral fusion with primary target B cells as established previously (Neuhierl et al., 2002). In addition, the assay unveils that virus stocks generated by ectopic expression of certain viral genes such as BBRF1 are not compromised with respect to virus fusion (**Fig. 8B**, right panel) but impaired in their infectivity (**Fig. 7B**). This observation shows that measuring virus fusion independent of virus infectivity is a valuable new parameter in the herpesvirus field. BBRF1 is a prime example, because it encodes the capsid portal protein (**Supplementary Table S1**) indispensable for loading EBV DNA into viral capsids. Upon BBRF1 knockdown, the virus stocks probably contain fewer infectious virions (they lack the viral DNA genome), but the virus stocks nevertheless show the identical fusogenic activity compared with wildtype virus indicating that fusion and *de novo* viral gene expression in EBV’s target cells are non-linked, independent steps in the multifaceted process of viral infection.

## Discussion

### The impact of individual EBV genes on virus yield and virus quality

The first recombinant EBV stocks were generated from a HEK293 cell line later termed 2089 EBV producer cells (Delecluse et al., 1998). EB virus supernatants were generated by transient transfection of an expression plasmid encoding BZLF1. We used a subclone of this original cell line for our analysis here, because it routinely yields up to 5×10^6^ infectious particles per ml with Raji cells used for the definition of infectious units. Despite this proven approach and reliable source of infectious EBV stocks, it has always been questionable, whether the HEK293 cell is the ideal host to propagate the virus, because this cell is probably of neuronal origin as it turned out later (Shaw et al., 2002). Regarding propagation of EBV a neuronal cell is an unusual cell type, which EBV does not infect. Thus, it seemed plausible to argue that HEK293 cells are compromised in yielding superior virus stocks because the cells are not expected to support efficient viral gene expression. Conversely, certain non-identified viral genes could exist that act as regulators of cellular or viral gene expression dampening or restricting virus production when expressed in an untimely fashion and/or at inadequate high levels in such a cell type. As the continuous search for other established cells that supports EBV production has not been successful in our laboratory, we invested into a systematic analysis of virus composition and quantity released from the 2089 EBV producer cell line. We asked whether ectopic expression of all available viral genes during virus synthesis in the producer cells might yield higher yields (indicative of an insufficient expression of this particular gene of interest) or if viral genes repress virus production, which could be an indication of a negative regulator of virus production. If true, the identification of such a viral regulatory function would help to design techniques to improve virus yield from HEK293 cells. Alternatively, the identification of a viral gene that boosts virus production during its enhanced expression in 2089 EBV producer cells will be another chance to improve virus yield.

In a single study from our lab in 2002 the function of a viral glycoprotein encoded by the EBV open reading frame BALF4 was investigated. Neuhierl et al. found that production of infectious EBV can be improved when BALF4 was ectopically expressed in the 2089 EBV producer cell line (Neuhierl et al., 2002). This finding, which was refined later (Neuhierl et al., 2009) indicated that BALF4 is a limiting gene product during EBV synthesis in HEK293 cells.

In our study here, we used this basic experimental design (with BALF4 as a proven positive control) and transfected BZLF1 together with 77 individual expression plasmids encoding single EBV genes listed in **Supplementary Table S1** into 2089 EBV producer cells. Supernatants from these experiments were investigated in four different analytic schemes as shown in **Figure 1**. The results from these analyses encompassing the parameters of physical particle concentration (**Fig. 3**), bioparticle concentration (**Fig. 5**) and virus titers (**Fig. 6**) all showed a good overall correlation (**Supplementary Fig. S4**). The fourth readout, a newly developed viral fusion assay using human primary B cells as targets adds an important aspect to our analysis (**Figure 8**).

Not unexpectedly, we found multiple viral genes which impacted virus functions and yield when ectopically expressed. Many different biological functions can have such a consequence. For example, high expression levels of viral genes can be toxic to cells or disturb regulatory circuits in the timely orchestrated steps of consecutive classes of viral genes (immediate early, early and late) that are characteristics of herpesvirus synthesis. Similarly, ectopic expression can cause problems in assembling virus components during morphogenesis as unbalanced levels could alter protein stoichiometry. Alternatively, viral genes could act as regulators of these steps and block virus synthesis when expressed at elevated levels or at an unsuitable time. To distinguish these possible scenarios, we invested into an efficient knockdown approach (**Supplementary Fig. S5**) with a panel of 75 shRNAs targeting 25 selected viral transcripts. Selection was based on a substantial reduction of virus titer and bioparticle concentration (’low bin’ genes in **Figures 5** and **6**) in line with a regulatory function. For comparison, we also chose genes that increased virus titer and bioparticle concentration. We failed to identify a viral gene with discrete regulatory function, because a knockdown of viral genes in the ‘low bin’ group of genes did not lead to virus titers or bioparticle concentrations beyond control levels as shown in the right panels of **Figure 7B and C**. Only the knockdown of a single gene, BFLF2, led to slightly increased virus titer (**Figure 7B**, left panel) but its ectopic expression also caused a very moderate increase, which argues against a regulatory function.

### ‘high bin’ and ‘low bin’ genes and their contribution to virus titer and bioparticle concentration

BALF4 scored best with regard to virus titers while the next ten top genes (BMRF1, BGRF1/BDRF1, LF1, BLRF3, BDLF2, BDLF3.5, BLLF2, BHLF1, BVLF1, BNLF2a) in **Figure 6** showed a moderate increase of virus titer, only. The role of BALF4 has been addressed previously by our laboratory and Richard Longnecker’s group (Herrold et al., 1996; Neuhierl et al., 2002) and more recently by Henri-Jacques Delecluse and colleagues (Neuhierl et al., 2009). Together, the literature agrees on the essential role of the BALF4 encoded gB glycoprotein, which is common to all herpesviruses and also called gp110 in the EBV field. The novel CD63:BlaM fusion assays presented in **Figure 8** documented the fusogenic characteristics of BALF4 to support the merger of the virus’ envelope with cell membranes probably within endosomes after receptor-mediated uptake of EBV (Chesnokova et al., 2015).

The known functions of the top ten genes **Figures 5** and **6** do not offer an immediate answer how they could contribute to the (mild) increase in virus titer. The genes fall into different functional categories and only BDLF2 and BILF1 belong to the class of membrane proteins similar to BALF4. BILF1 encodes a constitutively active G protein-coupled receptor presumably involved in signaling whereas BDLF2, which is one out of at least 11 EBV glycoprotein is a type II envelope protein with unknown functions (Gore and Hutt-Fletcher, 2009).

Down the gene list in **Figure 6**, ectopic expression of 16 viral genes reduced virus titers by half with some reducing it by a factor of 10 (BLRF1, BGLF5, BTRF1, BBRF1, BFRF1, BSLF2/BMLF1, BBRF3, BNRF1, BBLF1, BRLF1, BMRF2, BGLF4, BBRF2, BMLF1, LF2 and BALF1). Infection with EBV includes several discernable steps, i.e., virus binding, endocytosis, envelope fusion, viral transcription and translation of viral proteins. The Raji cell infection assay monitors all these steps leading to the expression of green fluorescence protein, which serves as a virally encoded surrogate marker and proxy of successful infection. Therefore, virus-cell binding, virus fusion, transcription and translation are indicators evaluating the quantity of infectious viruses in a given volume but also the quality of virus stocks. Whereas quantity is an obvious feature, quality is a rather vague parameter as it describes the virus fraction within a virus stock that is fully functional, i.e., infectious, whereas the remaining virus particles are not. This is, because they have not been properly assembled, they became damaged or essential virion components are underrepresented or entirely missing for various reasons. These particles are commonly called defective interfering particles, some of which we score as bioparticles. In our work their identification is solely based on the interaction of gp350 with CD21 on Elijah cells (**Fig. 4**). This was a surprising finding because gp350 is dispensable for infection of B cells (Janz et al., 2000) suggesting that other viral proteins might provide tethering functions as gp350 does. Our data shown in **Figure 4** clearly argue against this assumption.

### Groups of viral genes with comparable functionalities

Our systematic analysis of 77 viral genes did not identify a spectacular new function but revealed the contribution of many genes to EBV’s lytic phase including virus production and release. In an attempt to visualize the effects of the different genes tested we turned to spider charts (also called radar charts) to facilitate a comparison of six sets of quantitative data and to allow grouping of genes in classes according to their identified phenotypes.

The principal arrangement of 25 spider charts is shown in **Figure 9A**. Each spider chart covers four data sets when a gene is ectopically expressed and two data sets from shRNA knockdown experiments. Panel B shows the combined results obtained with BALF4. The effects are exceptional because its ectopic expression improves the fusogenic quality of virus stocks (**Fig. 8**) resulting in a much increased virus titer (**Fig. 6**) without a change in physical particle concentration (**Fig. 3**). Bioparticle concentration is reduced by half (**Figs. 5** and **8C**), but the virus titer is dramatically repressed upon knockdown of BALF4 (**Fig. 8B**) documenting that this viral gene encodes an essential virus function. No other EBV gene besides BALF4 shows such a profound phenotype confirming and extending previous data (Lee and Longnecker, 1997; Neuhierl et al., 2002; McShane and Longnecker, 2004; Neuhierl et al., 2009). The herpesvirus core fusion machinery consists of the conserved herpesvirus glycoproteins gB and gH/gL (Chandran and Hutt-Fletcher, 2007; Hutt-Fletcher, 2007) but, in contrast to BALF4 (which encodes gB) ectopic expression of gH or gL encoded by BXLF2 and BKRF2, respectively, did not reveal a clearly distinct phenotype in our model (**Figs. 3, 5, 6**) suggesting that BALF4 is expressed at insufficient levels in 2089 EBV producer cells. This interpretation could be problematic because we do not know BALF4 expression levels in virions from authentic virus producers such as B95.8 or Akata cells, for example. It is not within the scope of this work, but it would be interesting to express BALF4 ectopically also in these cells to investigate if gB is also limiting in virions from established EBV-infected B cell lines that give rise to progeny.

**Figure 9.**
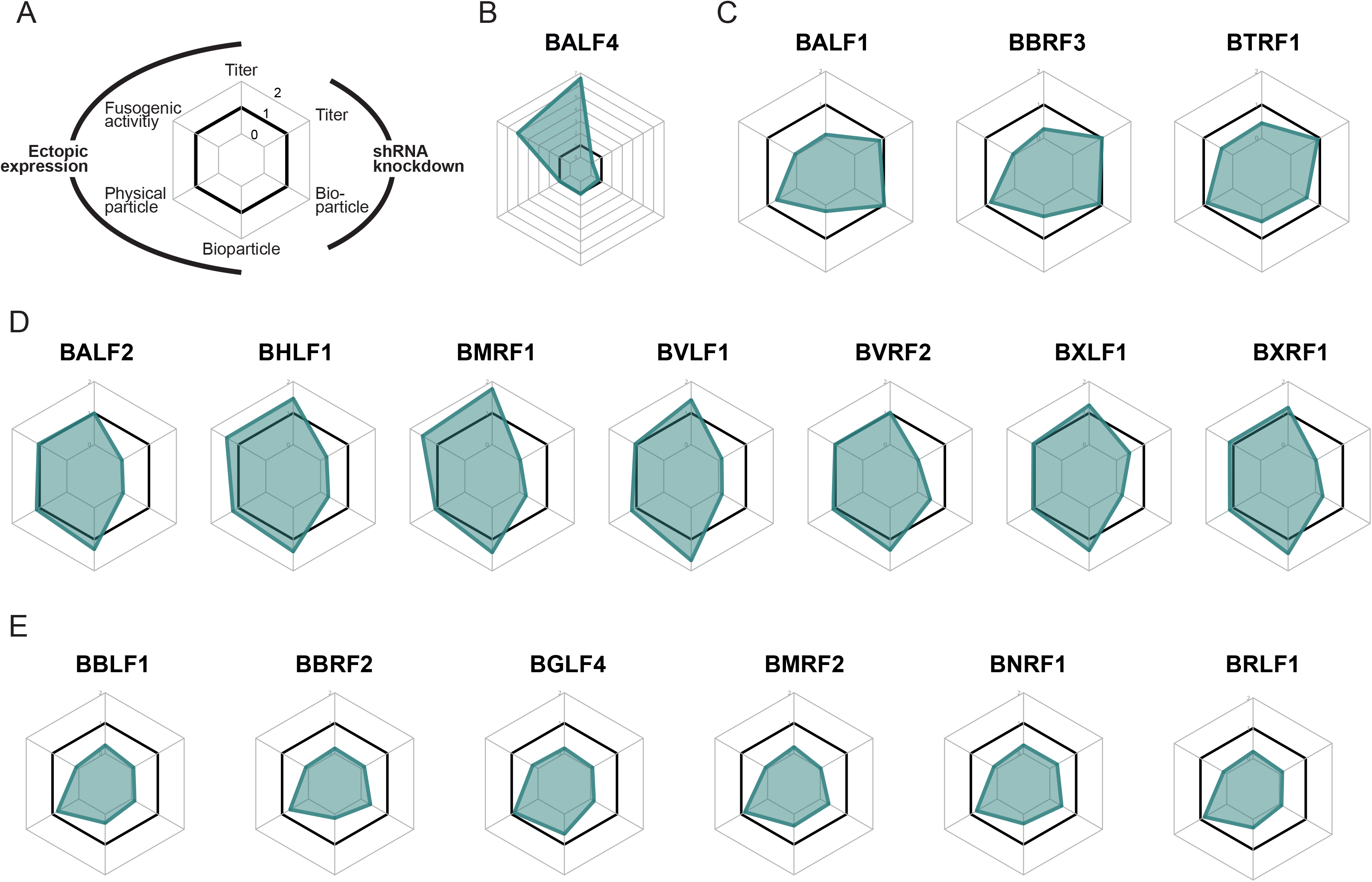
Spider charts visualizing the results of six parameters derived from the analysis of 17 selected viral genes. **(A)** Schematic view of the spider charts. Four of the six ‘legs’ of the diagram represent the calculated ratios of readouts encompassing ‘physical particle’, ‘bioparticle’, virus ‘titer’, and ‘fusogenic activity’ upon ectopic expression of selected viral genes. The remaining two legs show the consequences of shRNA mediated knockdown of a given gene affecting ‘virus ‘titer’ and ‘bioparticle’. Concentric hexameric rings indicate ratios of 1 (no change of the parameter), 0 (complete functional loss), and 2 (two-fold increase of the parameter). **(B)** The six ratios obtained with BALF4. Ectopic expression of BALF4 in 2089 EBV producer cells increases the virus titer and the fusogenic activity about seven- and four-fold, respectively. The shRNA mediated knockdown of BALF4 leads to an almost complete loss of the parameter virus ‘titer’ indicating that BALF4 is indispensable for virus synthesis. **(C)** Three viral genes, which, when expressed ectopically show reduced ratios of ‘bioparticles’, virus ‘titer’, and ‘fusogenic activity’. Their shRNA mediated knockdown does not affect ‘bioparticles’ and virus ‘titer’ ratios. **(D)** The selected seven viral genes have little effect on four parameters when expressed ectopically but their shRNA mediated knockdown severely compromised the parameter ratios ‘bioparticle’ and virus ‘titer’. **(E)** The parameter ratios of ‘bioparticle’, virus ‘titer’, and ‘fusogenic activity’ of six viral genes are compromised upon their ectopic expression and their shRNA mediated knockdown profoundly affects the parameters ‘bioparticle’ and virus ‘titer’ as well. Supplementary Figure S7 provides additional spider charts of other viral genes analyzed.

Spider charts with results from experimental testing of viral genes revealed three functional classes as shown in panels C, D and E of **Figure 9**. Panel C encompasses the genes BALF1, BBRF3 and BTRF1, which appear to be non-essential but, when ectopically expressed, compromise bioparticle release as well as virus titer and, probably as a consequence of both phenotypes, also the fusogenic activity of virus stocks. Very much in contrast, genes shown in panel D are essential as revealed by their shRNA mediated knockdown, but only some (BHLF1, BMRF1, BVLF1) improve virus titer and increase bioparticle concentration slightly upon ectopic expression. It thus seems as if this class of genes are optimally expressed upon lytic phase induction of 2089 EBV producer cells. Viral genes grouped in **Figure 9E** and **Supplementary Figure S7B** are similarly essential but their forced expression is detrimental indicating that they must be tightly regulated to support optimal virus production. It thus appears as if a certain level of gene expression is important during virus synthesis. A unique case seems to be BFLF2, which, upon its inactivation (Granato et al., 2008) as well as its induced ectopic expression (**Supplementary Fig. S7A**) was found to be crucial for virus production. Conversely, expression level of BFLF2 appears to be exceedingly high in 2089 EBV producer cells because virus titers increased in cells which express BFLF2 specific shRNAs (**Fig. 8B, Supplementary Fig. S7B**). Finally, the remaining four genes studied showed mixed phenotypes in **Supplementary Figure S7C**.

#### Achievements and obstacles

Our unbiased global attempt to identify viral genes with discrete functions in EBV’s lytic phase revealed a wide spectrum of phenotypes. Our study also suggests that the virus requires a close-to-perfect cellular host to support EBV production optimally. Contrary to our working hypothesis, HEK293 cells apparently fulfil most requirements EBV needs to escape from its latent phase and to give rise to progeny. Besides BALF4, the essential gB homologue present in all herpesviruses, other viral genes tested here did not stand out. It thus seems as if HEK293 cells provide an almost perfect cellular environment to support virus *de novo* synthesis.

A remaining mostly technical aspect is still worth discussing. From results presented in **Figure 2** the release of physical particles of viral nature can be estimated to be in the order of at least 5×10^8^ particles per ml using the NTA technology. This is a rough estimate at best because the background, which originates from extracellular vesicles contained in fetal bovine serum is already very high to begin with, and, in addition, non-induced HEK293 cells generate also at least 5×10^8^ physical particles per ml within three days of culture (**Fig. 2**, ‘conditioned Ex^−^’ medium). These observations make it very difficult to deduce the precise fraction of physical particles of viral nature given the abundance of all particles present. To address this, it will be prudent to design experiments with EBV releasing cells under conditions of serum-free, i.e. particle-free cell culture medium (**Fig. 2**). Moreover, one has to develop techniques to distinguish extracellular vesicles released from non-induced HEK293 cells in the culture from virions or virion-like vesicles that are decorated with or do contain viral components and are released from HEK293 cells that support EBV’s lytic phase. This fundamental aspect is certainly worth studying, but it requires more work to reveal the composition of particles released from 2089 EBV producer cells in comparison with established B cell lines upon their induction of EBV’s lytic phase.

## Materials and Methods

### Cell culture and cell lines

RPMI 1640 medium supplemented with 10 % FBS, 100 U/ml penicillin-streptomycin, 1 mM sodium pyruvate, 100 pM sodium selenite, and 0.04 % α-thioglycerols was used to culture all cells. 2089 carrying HEK293 EBV producer cells were cultivated in supplemented RPMI 1640 medium containing 100 μg/ml hygromycin B. EBV producer cells transduced with shRNA encoding lentiviral vectors were cultivated in supplemented RPMI cell culture medium and co-selected with 100 μg/ml hygromycin B and 3 μg/ml puromycin after initial selection with 10 μg/ml puromycin for 7 days. All cells were incubated in a 5 % CO_2_ and water-saturated atmosphere at 37°C.

### Transient transfection of 2089 EBV producer cells

6.5×10^5^ EBV producer cells (Delecluse et al., 1998) were seeded in 6-well cluster plates. After overnight incubation, the cell medium was exchanged with 2 ml EV depleted (Ex^−^) medium (**Figure 2**). 0.5 μg BZLF1 and 0.5 μg expression plasmid DNAs were mixed in a vial with 100 μl plain RPMI 1640. In another vial 100 μl plain RPMI 1640 was mixed with 6 μl PEI MAX (6 μl PEI MAX per 1 μg plasmid DNA). The content of both vials was combined rigorously mixed, the mixture was incubated at room temperature for 20 min and was added to the EBV producer cells in a single well of a 6-well cluster plate

### Assembly of ribonucleoprotein (RNP) complexes

Synthetic sgRNAs (Synthego) were dissolved in Nuclease-free Tris-EDTA Buffer (1× TE buffer) (Synthego) at a concentration of 100 μM. 6.5 μl of a 62 μM Cas9 V3 protein preparation (1081059; Integrated DNA Technologies) was added to 10 μl of the 100 μM gRNA solution. The mixture was diluted with sterile filtered (0.22 μm) PBS to a final volume of 50 μl and incubated at room temperature for 10 min to allow formation of RNPs at a final concentration of 8 μM Cas9. The two gRNAs specific for the *CR2* target gene were designed using the Synthego website. **Supplementary Table S5** provides the two gRNA sequences and their target nucleotide coordinates in exon 2 and exon 3 of the *CR2* locus on chromosome 1.

### Nucleofection of RNP complexes into Elijah cells

2×10^6^ Elijah cells were washed in PBS and resuspended in 20 μl P3 Primary Cell Nucleofector Solution buffer prepared with Supplement 1 buffer (Lonza) according to the manufacturer’s instructions (P3 Primary Cell 4D-Nucleofector X Kit S). For each of the two assembled gRNA-Cas9 complexes, 2.5 μl was mixed with the cell suspension, which was transferred to precooled (4 °C) well in a 16 well Nucleocuvette Strip (Lonza). Cells were nucleofected using the EH-100 program of Lonza’s protocol. 100 μl prewarmed (w/o supplements) RPMI 1640 medium was added to the cells, which were incubated for 15 min at 37 °C. The cells were transferred to a single well of a 24-well cluster plate and complete prewarmed cell culture medium containing 20 % FBS was added to a final volume of 220 μl to allow cell recovery. The cells were incubated at 37 °C, 5 % CO_2_.

### Flow cytometry and cell sorting

1×10^5^ Elijah cells were washed with and resuspended in 50 μl of FACS staining buffer (PBS, 1 % FBS, 2 mM EDTA) and stained with 1 μl of an APC-conjugated CD21 specific antibody (clone: HB5; 130-101-739, Miltenyi Biotec). The cells were incubated at 4 °C in the dark for 20 min and washed with and resuspended in 200 μl FACS staining buffer. Flow cytometry data were collected on a FACSCanto instrument (Becton Dickinson). For sorting, Elijah cells were stained with the CD21-APC antibody as described above. Cells were washed with and resuspended in the FACS staining buffer and filtered through a 100 μm mesh cell strainer to obtain single cell suspensions. Sorting was performed with a 100 μm wide nozzle, a velocity of about 8,000 events/second, a sorting mask of “4-way purity” using a BD FACS Aria IIIu instrument. The gating criteria included: (i) living cells, (ii) single cells, and (iii) CD21-negative cells. Data analysis was performed using the FlowJo software (version 10.4).

### shRNA expression vector construction and sequence design of shRNAs

The DNA sequences of selected EBV genes were obtained from the database of National Center for Biotechnology Information (NCBI). The FASTA format sequences were used for feeding the splashRNA tool (http://splashrna.mskcc.org) to predict potent shRNA sequences. According to the splashRNA score, the top three antisense guide sequences were chosen, ordered as synthetic oligonucleotides and inserted into the miR-3G frame of the basic shRNA construct (p6924 in our plasmid database). For each chosen EBV gene a knockdown pool of three shRNA constructs was designed and realized. The antisense guide sequences of individual EBV genes are marked in the entries shown in **Supplementary Table S2**, which have compatible ends to be cloned into AvrII and EcoRI restriction sites in the basic shRNA construct p6924. The shRNA constructs contain the puromycin resistance gene as selection marker.

### Virus titer measurement (infectivity)

The virus stocks were generated by transient DNA transfection of the 2089 EBV producer cells and the supernatants, harvested 3 days after transfection, were subsequently tested for infectious virus. Virus stocks were added to 1×10^5^ Raji cells in a volume of 2 ml cell culture medium and incubated for 3 days. The infected cells which express green fluorescence protein were determined and the fraction of GFP-positive cells, termed green Raji unit (GRU), was quantified by flow cytometry (BD FACSCanto).

### Bioparticle quantification (Elijah cell binding assay)

2×10^5^ Elijah cells were incubated with the harvested virus stocks. The mix was agitated on a mixing roller at 4 °C for 3 h. The cells were pelleted at 500 g at 4 °C for 10 min and washed with 1 ml ice-cold staining buffer (1% FBS and 2 mM EDTA in PBS). The anti-gp350 antibody (6G4) (Buschle et al., 2021) coupled to Alexa647 in 50 μl staining buffer (1:250 dilution) was added and the Elijah cells were incubated at 4 °C for 20 min. 1 ml staining buffer was added and the cells were washed and resuspended in 300 μl staining buffer and analyzed by flow cytometry (BD FACSCanto). The Elijah cells were analyzed for their fluorescence in the appropriate channel to obtain mean fluorescence intensity (MFI) data. MFI is an indirect measure of bound virus and correlates with the number of viral particles attached to the surface of Elijah cells.

### Physical particle measurement (Nanoparticle tracking analysis)

The physical particle concentration was measured using the ZetaView PMX 110 instrument (Particle Metrix), which can perform nanoparticle tracking analysis (NTA). Harvested virus stocks were diluted with PBS to adjust the concentration of particles to about 10^7^-10^8^ particles per ml. Standard calibration beads (102.7±1.3 nm) (Polysciences) were used to confirm the range of linearity of the instrument. 1 ml diluted supernatant samples were injected for analysis. All samples in the chamber were recorded at 11 positions in three cycles. Pre-acquisition parameters were set to 75 sensitivity, 70 shutter speed, a frame rate of 30 frames per second and 15 trace length. The post-acquisition parameters were set to a minimum brightness of 20, a minimum size of 5 pixels and a maximum size of 1000 pixels. Particle concentration and particle size were measured and documented and the images were analyzed using the ZetaView 8.04.02 software.

### Luciferase reporter construction

Three corresponding target/sense sequences were inserted into the dual luciferase reporter plasmid psiCHECK2 (p5264). The sequences were inserted downstream of the Renilla luciferase coding sequence using the restriction enzyme sites XhoI and NotI. Firefly luciferase is used as internal control. The shRNA target sequences are shown in **Supplementary Table S3**.

### Luciferase assay

2×10^5^ 293T were seeded in a 24-well cluster plate. After overnight incubation, cells were cotransfected with 100 ng psiCHECK2 luciferase reporter plasmid and 300 ng individual corresponding pCDH shRNA expression plasmids. An empty pCDH shRNA expression plasmid (p6924) was transfected as control. Cells were lysed after 24 h and luciferase activity was measured using the Orion luminometer (Berthold). To detect the knockdown efficiency of stably shRNA transduced 2089 EBV producer cells, 1×10^5^ producer cells were seeded in a 24-well cluster plate. After overnight incubation, cells were transfected with 100 ng corresponding psiCHECK2 luciferase reporter plasmid. An empty psiCHECK2 luciferase (p5264) reporter plasmid was transfected as control. After 24 h, cells were lysed and the luciferase activity was measured using the Orion luminometer.

### *ß*-lactamase (BlaM) fusion assay

EBV particles equipped with CD63:BlaM were generated by co-transfecting 6.5×10^5^ 2089 producer cells with expression plasmids encoding CD63:BlaM (p7200, 0.25 μg) and BZLF1 (p509, 0.25 μg) together with 0.5 μg plasmid DNA encoding single EBV genes to generate virus supernatants for further testing. The open reading frame of human CD63 is Cterminally fused to a codon-optimized **ß**-lactamase via a G_4_S linker and cloned into the expression plasmid pcDNA3.1 (+) (p5267). To evaluate the fusogenic activities of different virus stocks, 2×10^5^ human primary B cells were incubated with 5 μl virus supernatants for four hours. Cells were spun down and re-suspended in 100 μl of CCF4-AM staining solution in a 96-well plate. The staining solution consisted of 2 μl CCF4-AM (membrane-permeant ester forms of the negatively charged fluorescent **ß**-lactamase substrate), 8 μl Solution B (K1095, Thermo Fisher Scientific) and 10 μl of 250 mM Probenecid (P8761, Sigma) in 1 ml CO_2_-independent medium (18045-054, Thermo Fisher Scientific). After 16 h incubation in the dark at room temperature, the BlaM-positive recipient cells were analyzed by flow cytometry (BD LSRFortessa). The CCF4 FRET substrate is excited with 409 nm wavelength laser (violet). The non-cleaved substrate emits light with 520 nm wavelength (green) while the cleaved substrate emits light with a wavelength of 447 nm (blue).

### Isolation and preparation of human primary B cells from adenoids

Human primary B cells were purified from adenoidal tissues, which were chopped with blades and washed with PBS. The mashed tissues were filtered with 100 μm sterile strainer (352360, Falcon) and the cells were transferred to a sterile 50 ml tube. The volume was increased with PBS to 30 ml, 1 ml defibrinated sheep blood (SR0051D, Thermo Fisher Scientific) was added and mixed (to sediment T cells in the next step). The cell suspension was slowly layered on top of 15 ml Pancoll human (density: 1.077 g/ml) (P04-60500, PANBiotech) in a 50 ml tube. Samples were centrifuged at 1,900 rpm at room temperature for 30 min without brake. The interphase (white band) was collected and transferred to a new tube. The cells were washed three times with PBS and centrifuged at different speeds of 1,500, 1,400 and 1,200 rpm for 10 min.

### Physical particle concentration of formulated and conditioned cell culture media

Concentrations of physical particles in formulated and conditioned cell culture media were measured using a nanoparticle tracking analysis (NTA) instrument. Different formulated media and samples of cell culture media were analyzed. ‘RPMI^−^’ was plain RPMI 1640 medium purchased from the manufacturer; ‘RPMI^+^’ was plain RPMI 1640 medium with all supplements for cell culture excluding fetal bovine serum (FBS); ‘Ex^−^ medium’ was prepared by sedimenting 35 ml of ‘RPMI^+^’ medium supplemented with 20 % FBS by ultracentrifugation in a SW32 or SW28 swing-out rotor (Beckman) at 4 °C for at least 16 h. 30 ml of the supernatant was removed and diluted with an equal amount of ‘RPMI^+^’ medium. The medium was filtrated with 0.22 micron filter to yield 10 % FBS ‘Ex^−^ medium’. To quantitate particle numbers in conditioned cell medium, 2 ml of ‘Ex^−^ medium’ was incubated with noninduced or induced producer cells in 6-well cluster plates as shown in **Figure 2** for three days later, when the medium was collected, filtered using a 1.2 micron syringe filter and analyzed.

### Software tools

MacVector Version 18.0.0 (55) was used for in silico DNA cloning. FlowJo 10.4.2 was used for analysis and visualization of flow cytometry data. The ZetaView 8.04.02 software was used for nanoparticle tracking analysis. Prism 9 for macOS Version 9.0.1 (128) and R Version 4.0.5 with ggplot2 Version 3.3.3 and fmsb Version 0.7.0 packages were used for statistical analysis and visualization. Microsoft Word for Mac Version 16.45 and Microsoft Excel for Mac Version 16.45 were used for documentation and analysis. Adobe Illustrator 25.1 was used for composing figures. The diagrams were created with BioRender.com (https://biorender.com/).

## Acknowledgements

We thank our colleagues Regina Feederle and Reinhard Zeidler for the gift of the gp350 specific monoclonal antibody 6G4. This work was financially supported by grants from the Deutsche Forschungsgemeinschaft (grant number SFB1064/TP A13), Deutsche Krebshilfe (grant number 70112875) and German Center for Infection Research (grant number 07.814) to W.H.

## Author contributions

Y-FAC performed most experiments, DP provided technical and experimental support and advice. BM analyzed the data, calculated the statistics and designed the spider charts. EA performed the genomic knockout of CD21, Y-FAC and WH designed the scientific and experimental concept and wrote the manuscript. JM provided the expression library of EBV genes and their identities and Y-FAC, JM and WH finalized the manuscript.

---

**Supplementary Figure S1 Comparison of physical particle concentrations in virus samples generated by co-transfection of BZLF1 and single expression plasmids from a panel of 77 EBV genes including two controls.**

EB virus stocks were analyzed for their physical particle concentration by nanoparticle tracking analysis (NTA) as shown in Figure 3. EBV genes are marked according to their early or late expression characteristics as indicated. Mean and standard deviation of three biological replicates are shown. The horizontal lines indicate groups of 10 viral genes for better visualization.

**Supplementary Figure S2 Comparison of bioparticle concentrations in virus samples generated by BZLF1 transfection together with expression plasmids from a panel of 77 EBV genes and controls.**

Data are identical to those shown in Figure 5 but EBV genes are color-coded according to their early or late expression characteristics as indicated. The vertical lines provide 0.5- and 1.5-fold ratios. The horizontal lines indicate groups of 10 viral genes for better visualization.

**Supplementary Figure S3 Comparison of virus titers generated by co-transfection of BZLF1 together with expression plasmids from a panel of 77 EBV genes plus two controls.**

Data are identical to those shown in Figure 6 but EBV genes are marked according to their early or late expression characteristics. Two vertical lines indicate 0.5- and 1.5-fold ratios. The horizontal lines indicate groups of 10 viral genes for better visualization.

**Supplementary Figure S4 Statistical analysis of virus titer, bioparticle concentration, physical particle concentration, functional group category and expression time classification.**

**(A)** Heatmap of Spearman’s rank correlation coefficients between virus titer, bioparticle concentration and physical particle concentration measurements of all 77 individual virus supernatants is presented. Numbers in the grids indicate calculated Spearman correlation coefficients. For each measurement, a mean value from 3 replicates was considered prior to ranking and coefficient calculation. **(B)** Boxplots of virus titers (left), bioparticle concentrations (center) and physical particle concentrations (right) of all 77 individual virus supernatants, split by the functions of the viral gene co-expressed together with BZLF1 (DNA or capsid-associated proteins, membrane proteins, tegument proteins, non-structural proteins and unknown function). X-axes indicate the fold changes compared to the reference (Ctrl in Figures 3, 5, and 6). Each individual data point is the mean value of 3 individual measurements. The single red dot indicates the mean value of the virus supernatant obtained by co-transfection of BALF4 with BZLF1. **(C)** Boxplots of virus titers (left), bioparticle concentrations (center) and physical particle concentrations (right) of all 77 individual virus supernatants, split by the expression timing of the gene co-expressed together with BZLF1 (early, late, to be determined). X-axes indicate the fold changes compared to the reference (Ctrl in Figures 3, 5, and 6). The single red dot indicates the mean value of the virus supernatant of BALF4 co-expressed with BZLF1. An unpaired Wilcoxon rank sum test showed no significant differences, comparing the ‘early’ and ‘late’ groups. The median of the different populations is shown.

**Supplementary Figure S5 Functional evaluation of 75 shRNA constructs directed against 25 selected viral transcripts including controls.**

shRNA antisense guides for 25 EBV gene were generated (http://splashrna.mskcc.org) and three shRNA sequences per EBV target gene were individually cloned into the miR-3G frame of the basic shRNA vector (pCDH; p6924), which also encodes resistance against puromycin. A set of corresponding EBV target genes cloned into the psiCHECK2 dual luciferase reporter allows functional testing of the chosen shRNAs. **(A)** The flow chart depicts the experimental setup of the reporter assay shown in this panel. Transient reporter assays were conducted in 24-well cluster plates seeded with 2×10^5^ 293T cells, which were co-transfected with 25 individual reporter plasmids together with 25 single matching lentiviral shRNA expression plasmids. As reference and control, an empty pCDH lentiviral plasmid (p6924) was co-transfected with the indicated luciferase reporter (Ctrl). As additional controls, luciferase reporter plasmid with GFP and BALF4 as target genes were used in combination with three individual lentiviral vectors encoding GFP- and BALF4-specific shRNAs. After 24 h, luciferase activities were recorded and normalized to the reference. ‘High bin’ and ‘low bin’ samples are indicated. **(B)** The flow chart shows the experimental setup of introducing sets of three shRNA encoding lentiviruses into the 2089 EBV producer cell line and selection with puromycin. The individual cell lines stably transduced with three shRNA vectors each were analyzed by transient transfection with matching dual luciferase reporter plasmids encoding viral targets as in panel A. Mean and standard deviation of three biological replicates are shown. Asterisks indicate statistical significance as determined by using the unpaired two-tailed t test with Welch’s correction (**P≤ 0.01; ***P≤ 0.001). **(C)** The effect of three shRNAs stably introduced into the 2089 EBV producer cell line and directed against GFP was analyzed by flow cytometry. GFP-negative cells were used as control. GFP expression of parental 2089 EBV producer cells (2089) and a derivative stably transduced with an empty pCDH lentiviral plasmid (p6924) as an shRNA control (shCtrl) are shown. The x-axis denotes intensity of GFP fluorescence.

**Supplementary Figure S6 Comparison of the infectivity of engineered EBV stocks generated from 2089 EBV producer cells after dual transfection of BZLF1 and single expression plasmids or after triple transfection with BZLF1, single expression plasmids and CD63:BlaM.**

Supernatants analyzed in panel B of Figure 8 were tested for their infectivity using Raji cells and GFP expression as indicator of viral infection as schematically shown in panel A of Figure 8. For comparison, the 2089 EBV producer cell line was transiently transfected with 25 individual expression plasmids encoding viral genes of the ‘high bin’, ‘low bin’ groups and three ‘random’ genes together with BZLF1 p509, only, omitting the CD63:*ß*-lactamase (CD63:BlaM) expression plasmid p7200. The pairwise comparison of supernatants generated with and without CD63:BlaM is shown.

**Supplementary Figure S7 Additional spider charts visualizing the results of six parameters derived from the analysis of selected viral genes.**

Shown are additional charts with parameter ratios obtained from 8 viral genes supplementing data shown in Figure 9.

