## Supplementary figures for "Systematic analysis of Epstein-Barr virus genes and their individual contribution to virus production and composition"

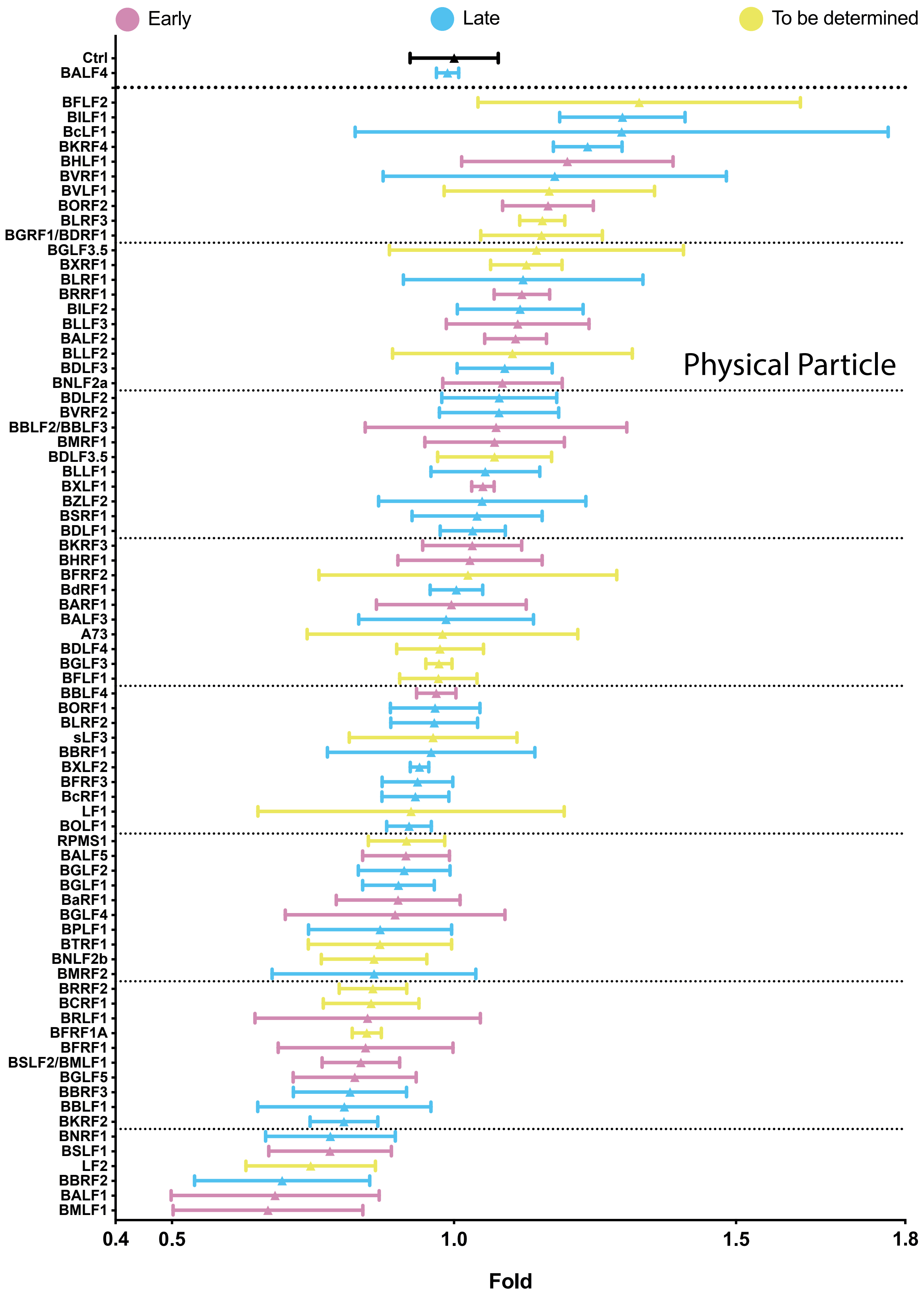


Supplementary Figure S1

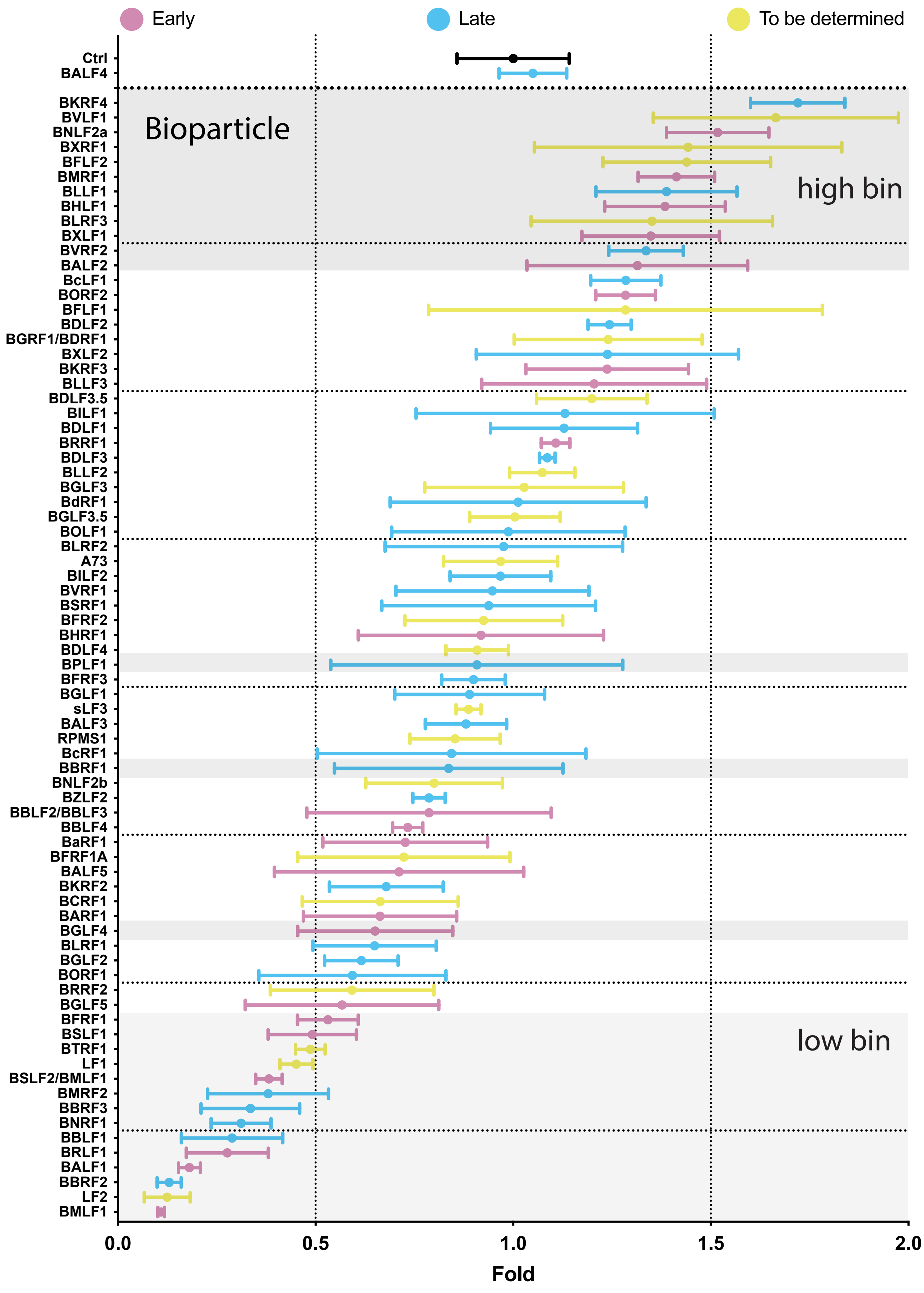


Supplementary Figure S2

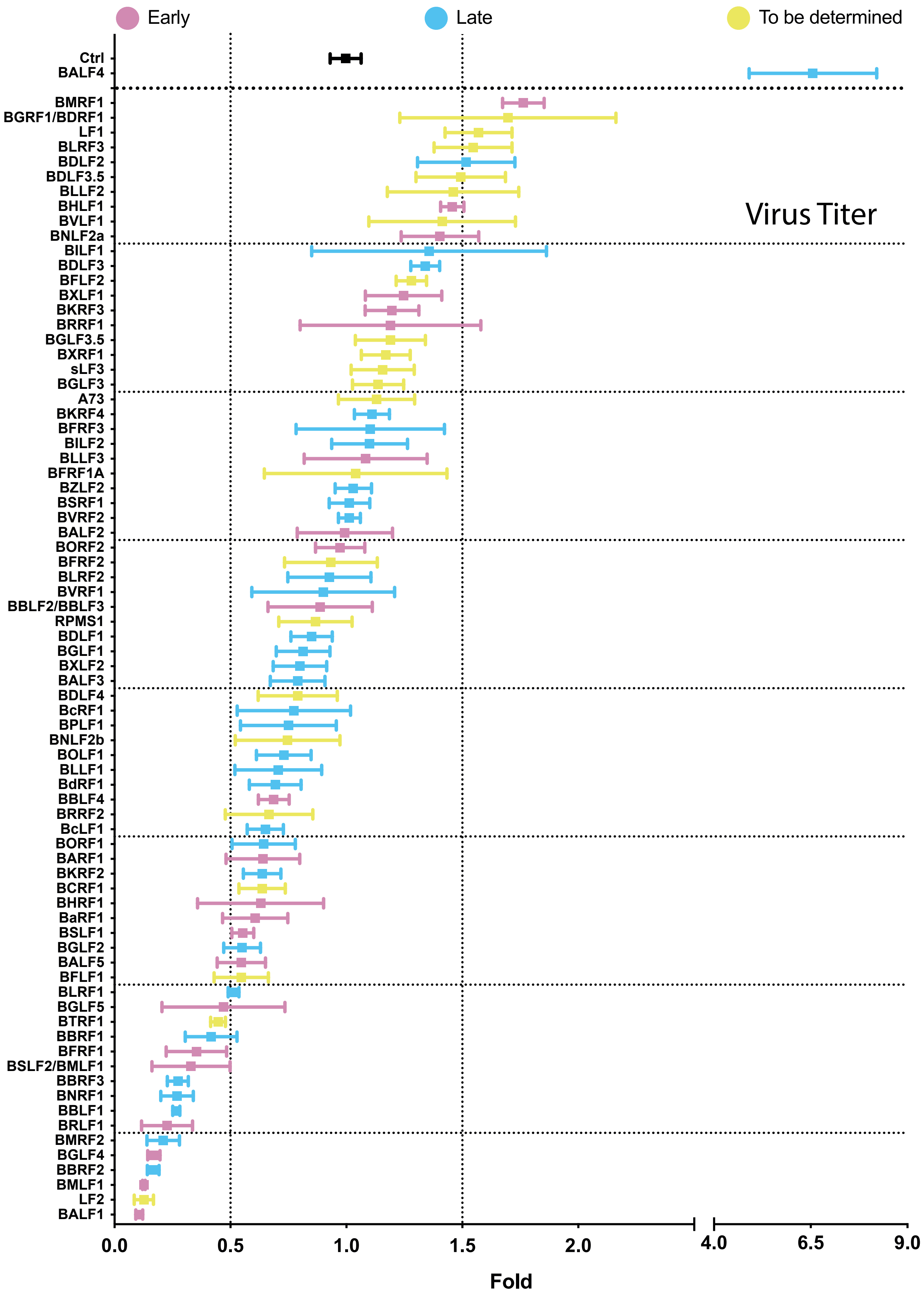


Supplementary Figure S3

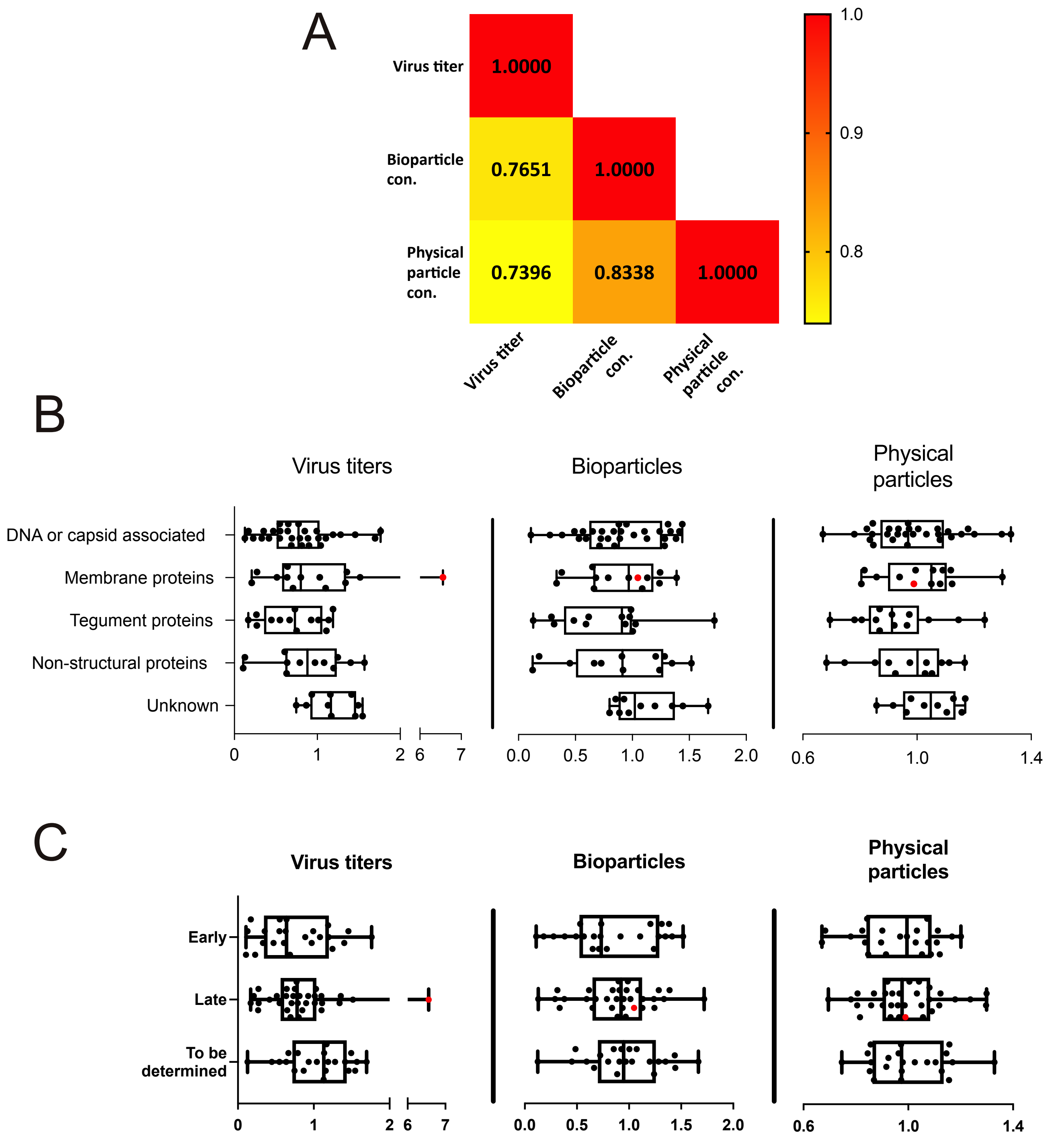


Supplementary Figure S4

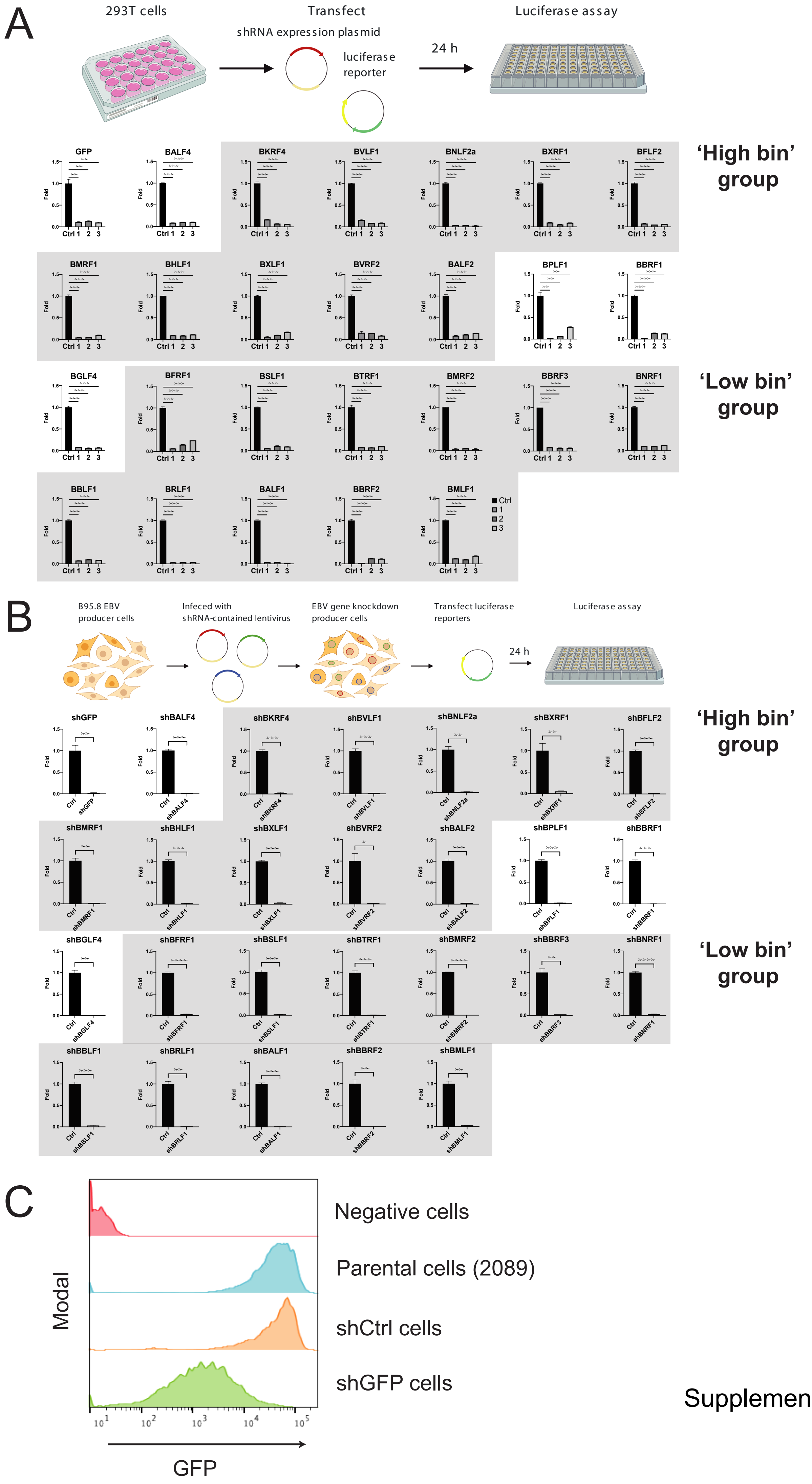


Supplementary Figure S5

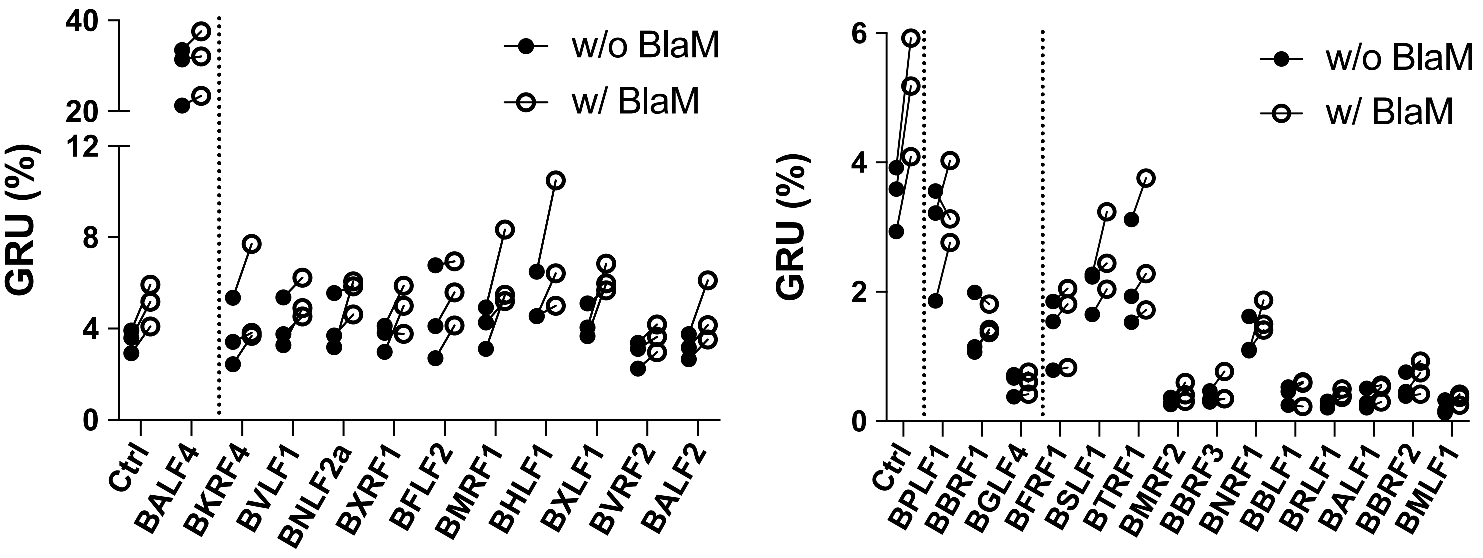


Supplementary Figure S6

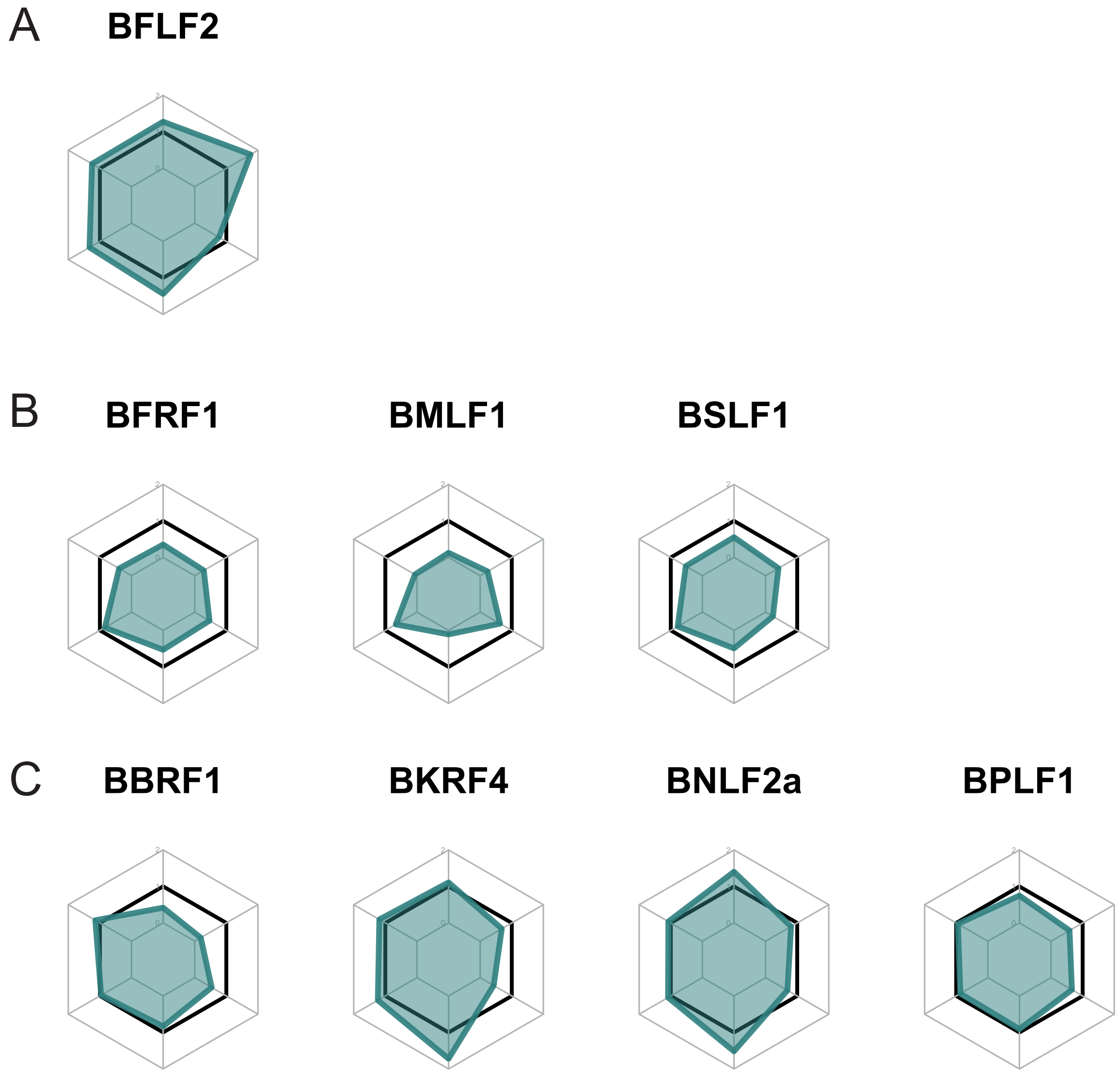


Supplementary Figure S7
