## Supplementary Table S1 for "Systematic analysis of Epstein-Barr virus genes and their individual contribution to virus production and composition"

**Table S1. Overview of identity, related function and expression timing of 78 EBV genes**

| Gene | Identity | Related Function | Early/Late phase^1^ |
| --- | --- | --- | --- |
| A73 | Protein A73 | Unknown | TBD |
| BALF1 | Apoptosis regulator BALF1 | Non-structural protein | E |
| BALF2 | Single-stranded DNA-binding protein | DNA or Capsid associated | E |
| BALF3 | DNA packaging terminase subunit 2 | DNA or Capsid associated | L |
| BALF4 | Envelope glycoprotein B, gB | Membrane protein | L |
| BALF5 | DNA polymerase catalytic subunit | DNA or Capsid associated | E |
| BARF1 | Homolog of the human proto-oncogene c-fms | Membrane protein | E |
| BaRF1 | Ribonucleotide reductase subunit 2 | Non-structural protein | E |
| BBLF1 | Myristylated tegument protein | Tegument | L |
| BBLF2/BBLF3 | Helicase-primase subunit | DNA or Capsid associated | E |
| BBLF4 | Helicase-primase helicase subunit | DNA or Capsid associated | E |
| BBRF1 | Capsid portal protein | DNA or Capsid associated | L |
| BBRF2 | Tegument protein UL7 | Tegument | L |
| BBRF3 | Envelope glycoprotein M | Membrane protein | L |
| BcLF1 | Major capsid protein | DNA or Capsid associated | L |
| BCRF1 | Interleukin-10 | Non-structural protein | TBD |
| BcRF1 | Viral TATA box binding protein | DNA or Capsid associated | L |
| BDLF1 | Capsid triplex subunit 2 | DNA or Capsid associated | L |
| BDLF2 | Envelope glycoprotein 48 | Membrane protein | L |
| BDLF3 | Envelope glycoprotein 150 | Membrane protein | L |
| BDLF3.5 | Protein UL91 | Unknown | TBD |
| BDLF4 | Protein UL92 | Non-structural protein | TBD |
| BdRF1 | Capsid scaffold protein | DNA or Capsid associated | L |
| BFLF1 | DNA packaging protein UL32 | DNA or Capsid associated | TBD |
| BFLF2 | Nuclear egress lamina protein | DNA or Capsid associated | TBD |
| BFRF1 | Nuclear egress membrane protein | DNA or Capsid associated | E |
| BFRF1A | DNA packaging protein UL33 | DNA or Capsid associated | TBD |
| BFRF2 | Protein UL49 | Unknown | TBD |
| BFRF3 | Small capsid protein | DNA or Capsid associated | L |
| BGLF1 | DNA packaging tegument protein UL17 | DNA or Capsid associated | L |
| BGLF2 | Tegument protein UL16 | Tegument | L |
| BGLF3 | Protein UL95 | Tegument | TBD |
| BGLF3.5 | Tegument protein UL14 | Tegument | TBD |
| BGLF4 | Tegument serine/threonine protein kinase | DNA or Capsid associated | E |
| BGLF5 | Deoxyribonuclease | DNA or Capsid associated | E |
| BGRF1/BDRF1 | DNA packaging terminase subunit 1 | DNA or Capsid associated | TBD |
| BHLF1 | Protein BHLF1 | DNA or Capsid associated | E |
| BHRF1 | Apoptosis regulator BHRF1 | Non-structural protein | E |
| BILF1 | Membrane protein BILF1 | Membrane protein | L |
| BILF2 | Membrane protein BILF2 | Membrane protein | L |
| BKRF2 | Envelope glycoprotein L, gL | Membrane protein | L |
| BKRF3 | Uracil-DNA glycosylase | Non-structural protein | E |
| BKRF4 | Tegument protein G45 | Tegument | L |
| BLLF1 | Glycoprotein 350 | Membrane protein | L |
| BLLF2 | Protein BLLF2 | Unknown | TBD |
| BLLF3 | Deoxyuridine triphosphatase | Non-structural protein | E |
| BLRF1 | Envelope glycoprotein N | Membrane protein | L |
| BLRF2 | Virion protein G52 | Tegument | L |
| BLRF3 | Unknown | Unknown | TBD |
| BMLF1 | EB2 | DNA or Capsid associated | E |
| BMRF1 | DNA polymerase processivity subunit | DNA or Capsid associated | E |
| BMRF2 | Envelope protein UL43 | Membrane protein | L |
| BNLF2a | Protein BNLF2a | Non-structural protein | E |
| BNLF2b | Protein BNLF2b | Unknown | TBD |
| BNRF1 | Tegument protein G75 | Tegument | L |
| BOLF1 | Tegument protein UL37 | Tegument | L |
| BORF1 | Capsid triplex subunit 1 | DNA or Capsid associated | L |
| BORF2 | Ribonucleotide reductase subunit 1 | Non-structural protein | E |
| BPLF1 | Large tegument protein | Tegument | L |
| BRLF1 | Protein Rta | DNA or Capsid associated | IE |
| BRRF1 | Protein G49 | DNA or Capsid associated | E |
| BRRF2 | Tegument protein G48 | Tegument | TBD |
| BSLF1 | Helicase-primase primase subunit | DNA or Capsid associated | E |
| BSLF2/BMLF1 | Multifunctional expression regulator | DNA or Capsid associated | E |
| BSRF1 | Tegument protein UL51 | Tegument | L |
| BTRF1 | Tegument protein UL88 | Tegument | TBD |
| BVLF1 | Protein UL79 | Unknown | TBD |
| BVRF1 | DNA packaging tegument protein UL25 | DNA or Capsid associated | L |
| BVRF2 | Capsid maturation protease | DNA or Capsid associated | L |
| BXLF1 | Thymidine kinase | Non-structural protein | E |
| BXLF2 | Envelope glycoprotein H, gH | Membrane protein | L |
| BXRF1 | Nuclear protein UL24 | Unknown | TBD |
| BZLF1 | Protein Zta | DNA or Capsid associated | IE |
| BZLF2 | Envelope glycoprotein 42 | Membrane protein | L |
| LF1 | Protein G10 | Non-structural protein | TBD |
| LF2 | Virion protein G11 | Non-structural protein | TBD |
| RPMS1 | Protein RPMS1 | Unknown | TBD |
| sLF3 | Protein LF3 | Unknown | TBD |

1: E, early; IE, immediate early; L, late; TBD: to be determined.

Table was adapted according to Johannsen et al. Proteins of purified Epstein-Barr virus. PNAS, 2004; Arvin A et al. Human Herpesviruses: Biology, Therapy, and Immunoprophylaxis. Cambridge: Cambridge University Press; 2007. PMID: 21348071; Cai et al. Characterization of the subcellular localization of Epstein-Barr virus encoded proteins in live cells. Oncotarget, 2017.
