## Supplementary Table S2 for "Systematic analysis of Epstein-Barr virus genes and their individual contribution to virus production and composition"

**Table S2. Overview of synthetic oligonucleotides cloned into the basic lentiviral shRNA expression vector plasmid p6924 to be stably introduced into 2089 EBV producer cells**

| Target gene (plasmid #) | Oligonucleotide |
| --- | --- |
| BALF4 (p6515) | shRNA 6954 for  CTAGGTTTATGTTTGGATGAACTGACATACGCGTATCCGTCTAATGTTGTCTTTAAACACCATGTAGTGAAATATATATTAAACATGGTGTTTAAAGACAACATTTTACGGTAACGCGG |
|  | shRNA 6954 rev  AATTCCGCGTTACCGTAAAATGTTGTCTTTAAACACCATGTTTAATATATATTTCACTACATGGTGTTTAAAGACAACATTAGACGGATACGCGTATGTCAGTTCATCCAAACATAAAC |
|  | shRNA 6955 for  CTAGGTTTATGTTTGGATGAACTGACATACGCGTATCCGTCTTTAGTGAGATGAAGGTCTGCAGTAGTGAAATATATATTAAACTGCAGACCTTCATCTCACTAATTACGGTAACGCGG |
|  | shRNA 6955 rev  AATTCCGCGTTACCGTAATTAGTGAGATGAAGGTCTGCAGTTTAATATATATTTCACTACTGCAGACCTTCATCTCACTAAAGACGGATACGCGTATGTCAGTTCATCCAAACATAAAC |
|  | shRNA 6956 for  CTAGGTTTATGTTTGGATGAACTGACATACGCGTATCCGTCTTATAAAATATGTAATGGCTTCGTAGTGAAATATATATTAAACGAAGCCATTACATATTTTATATTACGGTAACGCGG |
|  | shRNA 6956 rev  AATTCCGCGTTACCGTAATATAAAATATGTAATGGCTTCGTTTAATATATATTTCACTACGAAGCCATTACATATTTTATAAGACGGATACGCGTATGTCAGTTCATCCAAACATAAAC |
| BKRF4 (p6562) | shRNA 7296 top  CTAGGTTTATGTTTGGATGAACTGACATACGCGTATCCGTCTGTGTTTATTGTATGTATTGGGGTAGTGAAATATATATTAAACCCCAATACATACAATAAACACTTACGGTAACGCGG |
|  | shRNA 7296 bot  AATTCCGCGTTACCGTAAGTGTTTATTGTATGTATTGGGGTTTAATATATATTTCACTACCCCAATACATACAATAAACACAGACGGATACGCGTATGTCAGTTCATCCAAACATAAAC |
|  | shRNA 7297 top  CTAGGTTTATGTTTGGATGAACTGACATACGCGTATCCGTCTTTATTGTATGTATTGGGACTTGTAGTGAAATATATATTAAACAAGTCCCAATACATACAATAATTACGGTAACGCGG |
|  | shRNA 7297 bot  AATTCCGCGTTACCGTAATTATTGTATGTATTGGGACTTGTTTAATATATATTTCACTACAAGTCCCAATACATACAATAAAGACGGATACGCGTATGTCAGTTCATCCAAACATAAAC |
|  | shRNA 7298 top  CTAGGTTTATGTTTGGATGAACTGACATACGCGTATCCGTCTATTGTATGTATTGGGACTTGAGTAGTGAAATATATATTAAACTCAAGTCCCAATACATACAATTTACGGTAACGCGG |
|  | shRNA 7298 bot  AATTCCGCGTTACCGTAAATTGTATGTATTGGGACTTGAGTTTAATATATATTTCACTACTCAAGTCCCAATACATACAATAGACGGATACGCGTATGTCAGTTCATCCAAACATAAAC |
| BVLF1 (p6846) | shRNA 7299 top  CTAGGTTTATGTTTGGATGAACTGACATACGCGTATCCGTCTAAATAAACTCATCGCACGGGGGTAGTGAAATATATATTAAACCCCCGTGCGATGAGTTTATTTTTACGGTAACGCGG |
|  | shRNA 7299 bot  AATTCCGCGTTACCGTAAAAATAAACTCATCGCACGGGGGTTTAATATATATTTCACTACCCCCGTGCGATGAGTTTATTTAGACGGATACGCGTATGTCAGTTCATCCAAACATAAAC |
|  | shRNA 7300 top  CTAGGTTTATGTTTGGATGAACTGACATACGCGTATCCGTCTTCTTTAGCATCTTCAGGAGGAGTAGTGAAATATATATTAAACTCCTCCTGAAGATGCTAAAGATTACGGTAACGCGG |
|  | shRNA 7300 bot  AATTCCGCGTTACCGTAATCTTTAGCATCTTCAGGAGGAGTTTAATATATATTTCACTACTCCTCCTGAAGATGCTAAAGAAGACGGATACGCGTATGTCAGTTCATCCAAACATAAAC |
|  | shRNA 7301 top  CTAGGTTTATGTTTGGATGAACTGACATACGCGTATCCGTCTAAAAGTTAACACTTAGGGTCAGTAGTGAAATATATATTAAACTGACCCTAAGTGTTAACTTTTTTACGGTAACGCGG |
|  | shRNA 7301 bot  AATTCCGCGTTACCGTAAAAAAGTTAACACTTAGGGTCAGTTTAATATATATTTCACTACTGACCCTAAGTGTTAACTTTTAGACGGATACGCGTATGTCAGTTCATCCAAACATAAAC |
| BNLF2a (p6839) | shRNA 7302 top  CTAGGTTTATGTTTGGATGAACTGACATACGCGTATCCGTCTTTATTTATTGCATCACAAGTCGTAGTGAAATATATATTAAACGACTTGTGATGCAATAAATAATTACGGTAACGCGG |
|  | shRNA 7302 bot  AATTCCGCGTTACCGTAATTATTTATTGCATCACAAGTCGTTTAATATATATTTCACTACGACTTGTGATGCAATAAATAAAGACGGATACGCGTATGTCAGTTCATCCAAACATAAAC |
|  | shRNA 7303 top  CTAGGTTTATGTTTGGATGAACTGACATACGCGTATCCGTCTTATTTATTGCATCACAAGTCAGTAGTGAAATATATATTAAACTGACTTGTGATGCAATAAATATTACGGTAACGCGG |
|  | shRNA 7303 bot  AATTCCGCGTTACCGTAATATTTATTGCATCACAAGTCAGTTTAATATATATTTCACTACTGACTTGTGATGCAATAAATAAGACGGATACGCGTATGTCAGTTCATCCAAACATAAAC |
|  | shRNA 7304 top  CTAGGTTTATGTTTGGATGAACTGACATACGCGTATCCGTCTTTATTGCATCACAAGTCACATGTAGTGAAATATATATTAAACATGTGACTTGTGATGCAATAATTACGGTAACGCGG |
|  | shRNA 7304 bot  AATTCCGCGTTACCGTAATTATTGCATCACAAGTCACATGTTTAATATATATTTCACTACATGTGACTTGTGATGCAATAAAGACGGATACGCGTATGTCAGTTCATCCAAACATAAAC |
| BXRF1 (p6848) | shRNA 7305 top  CTAGGTTTATGTTTGGATGAACTGACATACGCGTATCCGTCTTGTAAATCTTATTGTCCGCGAGTAGTGAAATATATATTAAACTCGCGGACAATAAGATTTACATTACGGTAACGCGG |
|  | shRNA 7305 bot  AATTCCGCGTTACCGTAATGTAAATCTTATTGTCCGCGAGTTTAATATATATTTCACTACTCGCGGACAATAAGATTTACAAGACGGATACGCGTATGTCAGTTCATCCAAACATAAAC |
|  | shRNA 7306 top  CTAGGTTTATGTTTGGATGAACTGACATACGCGTATCCGTCTTAAATTCTACAATATAACACCGTAGTGAAATATATATTAAACGGTGTTATATTGTAGAATTTATTACGGTAACGCGG |
|  | shRNA 7306 bot  AATTCCGCGTTACCGTAATAAATTCTACAATATAACACCGTTTAATATATATTTCACTACGGTGTTATATTGTAGAATTTAAGACGGATACGCGTATGTCAGTTCATCCAAACATAAAC |
|  | shRNA 7307 top  CTAGGTTTATGTTTGGATGAACTGACATACGCGTATCCGTCTTGATAATCTCAAAGAGGGTGTGTAGTGAAATATATATTAAACACACCCTCTTTGAGATTATCATTACGGTAACGCGG |
|  | shRNA 7307 bot  AATTCCGCGTTACCGTAATGATAATCTCAAAGAGGGTGTGTTTAATATATATTTCACTACACACCCTCTTTGAGATTATCAAGACGGATACGCGTATGTCAGTTCATCCAAACATAAAC |
| BFLF2 (p6644) | shRNA 7308 top  CTAGGTTTATGTTTGGATGAACTGACATACGCGTATCCGTCTTATTTTCCAAAATGAGCTGGGGTAGTGAAATATATATTAAACCCCAGCTCATTTTGGAAAATATTACGGTAACGCGG |
|  | shRNA 7308 bot  AATTCCGCGTTACCGTAATATTTTCCAAAATGAGCTGGGGTTTAATATATATTTCACTACCCCAGCTCATTTTGGAAAATAAGACGGATACGCGTATGTCAGTTCATCCAAACATAAAC |
|  | shRNA 7309 top  CTAGGTTTATGTTTGGATGAACTGACATACGCGTATCCGTCTTTATTTTCCAAAATGAGCTGGGTAGTGAAATATATATTAAACCCAGCTCATTTTGGAAAATAATTACGGTAACGCGG |
|  | shRNA 7309 bot  AATTCCGCGTTACCGTAATTATTTTCCAAAATGAGCTGGGTTTAATATATATTTCACTACCCAGCTCATTTTGGAAAATAAAGACGGATACGCGTATGTCAGTTCATCCAAACATAAAC |
|  | shRNA 7310 top  CTAGGTTTATGTTTGGATGAACTGACATACGCGTATCCGTCTTTGATAGGACTGTACCAGGTCGTAGTGAAATATATATTAAACGACCTGGTACAGTCCTATCAATTACGGTAACGCGG |
|  | shRNA 7310 bot  AATTCCGCGTTACCGTAATTGATAGGACTGTACCAGGTCGTTTAATATATATTTCACTACGACCTGGTACAGTCCTATCAAAGACGGATACGCGTATGTCAGTTCATCCAAACATAAAC |
| BMRF1 (p5106) | shRNA 7311 top (same as 6963 for)  CTAGGTTTATGTTTGGATGAACTGACATACGCGTATCCGTCTAGGATTTAATGAATGTCACCAGTAGTGAAATATATATTAAACTGGTGACATTCATTAAATCCTTTACGGTAACGCGG |
|  | shRNA 7311 bot (same as 6963 rev)  AATTCCGCGTTACCGTAAAGGATTTAATGAATGTCACCAGTTTAATATATATTTCACTACTGGTGACATTCATTAAATCCTAGACGGATACGCGTATGTCAGTTCATCCAAACATAAAC |
|  | shRNA 7312 top  CTAGGTTTATGTTTGGATGAACTGACATACGCGTATCCGTCTAAAATAACACTAAGATCCAACGTAGTGAAATATATATTAAACGTTGGATCTTAGTGTTATTTTTTACGGTAACGCGG |
|  | shRNA 7312 bot  AATTCCGCGTTACCGTAAAAAATAACACTAAGATCCAACGTTTAATATATATTTCACTACGTTGGATCTTAGTGTTATTTTAGACGGATACGCGTATGTCAGTTCATCCAAACATAAAC |
|  | shRNA 7313 top  CTAGGTTTATGTTTGGATGAACTGACATACGCGTATCCGTCTTATCATCATATTCCATAGTGAGTAGTGAAATATATATTAAACTCACTATGGAATATGATGATATTACGGTAACGCGG |
|  | shRNA 7313 bot  AATTCCGCGTTACCGTAATATCATCATATTCCATAGTGAGTTTAATATATATTTCACTACTCACTATGGAATATGATGATAAGACGGATACGCGTATGTCAGTTCATCCAAACATAAAC |
| BHLF1 (p7080) | shRNA 7314 top  CTAGGTTTATGTTTGGATGAACTGACATACGCGTATCCGTCTTAAACAGTTTATTGATAGGTGGTAGTGAAATATATATTAAACCACCTATCAATAAACTGTTTATTACGGTAACGCGG |
|  | shRNA 7314 bot  AATTCCGCGTTACCGTAATAAACAGTTTATTGATAGGTGGTTTAATATATATTTCACTACCACCTATCAATAAACTGTTTAAGACGGATACGCGTATGTCAGTTCATCCAAACATAAAC |
|  | shRNA 7315 top  CTAGGTTTATGTTTGGATGAACTGACATACGCGTATCCGTCTTGACATGTAGGTGAGTAGTGTGTAGTGAAATATATATTAAACACACTACTCACCTACATGTCATTACGGTAACGCGG |
|  | shRNA 7315 bot  AATTCCGCGTTACCGTAATGACATGTAGGTGAGTAGTGTGTTTAATATATATTTCACTACACACTACTCACCTACATGTCAAGACGGATACGCGTATGTCAGTTCATCCAAACATAAAC |
|  | shRNA 7316 top  CTAGGTTTATGTTTGGATGAACTGACATACGCGTATCCGTCTTACTTTTAGAGTGTAGTGTACGTAGTGAAATATATATTAAACGTACACTACACTCTAAAAGTATTACGGTAACGCGG |
|  | shRNA 7316 bot  AATTCCGCGTTACCGTAATACTTTTAGAGTGTAGTGTACGTTTAATATATATTTCACTACGTACACTACACTCTAAAAGTAAGACGGATACGCGTATGTCAGTTCATCCAAACATAAAC |
| BXLF1 (p6571) | shRNA 7320 top  CTAGGTTTATGTTTGGATGAACTGACATACGCGTATCCGTCTTAAGTACACTAAAGATGCTGTGTAGTGAAATATATATTAAACACAGCATCTTTAGTGTACTTATTACGGTAACGCGG |
|  | shRNA 7320 bot  AATTCCGCGTTACCGTAATAAGTACACTAAAGATGCTGTGTTTAATATATATTTCACTACACAGCATCTTTAGTGTACTTAAGACGGATACGCGTATGTCAGTTCATCCAAACATAAAC |
|  | shRNA 7321 top  CTAGGTTTATGTTTGGATGAACTGACATACGCGTATCCGTCTTGTAAATTAACTTGTAGCGGTGTAGTGAAATATATATTAAACACCGCTACAAGTTAATTTACATTACGGTAACGCGG |
|  | shRNA 7321 bot  AATTCCGCGTTACCGTAATGTAAATTAACTTGTAGCGGTGTTTAATATATATTTCACTACACCGCTACAAGTTAATTTACAAGACGGATACGCGTATGTCAGTTCATCCAAACATAAAC |
|  | shRNA 7322 top  CTAGGTTTATGTTTGGATGAACTGACATACGCGTATCCGTCTAACTTACTAAACTCGCGCCCAGTAGTGAAATATATATTAAACTGGGCGCGAGTTTAGTAAGTTTTACGGTAACGCGG |
|  | shRNA 7322 bot  AATTCCGCGTTACCGTAAAACTTACTAAACTCGCGCCCAGTTTAATATATATTTCACTACTGGGCGCGAGTTTAGTAAGTTAGACGGATACGCGTATGTCAGTTCATCCAAACATAAAC |
| BVRF2 (p6847) | shRNA 7264 top  CTAGGTTTATGTTTGGATGAACTGACATACGCGTATCCGTCTTTAAATACGACTCGGCTGGGAGTAGTGAAATATATATTAAACTCCCAGCCGAGTCGTATTTAATTACGGTAACGCGG |
|  | shRNA 7264 bot  AATTCCGCGTTACCGTAATTAAATACGACTCGGCTGGGAGTTTAATATATATTTCACTACTCCCAGCCGAGTCGTATTTAAAGACGGATACGCGTATGTCAGTTCATCCAAACATAAAC |
|  | shRNA 7265top  CTAGGTTTATGTTTGGATGAACTGACATACGCGTATCCGTCTAAGTGTTAACTTTTACCTGTGGTAGTGAAATATATATTAAACCACAGGTAAAAGTTAACACTTTTACGGTAACGCGG |
|  | shRNA 7265 bot  AATTCCGCGTTACCGTAAAAGTGTTAACTTTTACCTGTGGTTTAATATATATTTCACTACCACAGGTAAAAGTTAACACTTAGACGGATACGCGTATGTCAGTTCATCCAAACATAAAC |
|  | shRNA 7266 top  CTAGGTTTATGTTTGGATGAACTGACATACGCGTATCCGTCTTTGATGAGGCTGAAATCCGTAGTAGTGAAATATATATTAAACTACGGATTTCAGCCTCATCAATTACGGTAACGCGG |
|  | shRNA 7266 bot  AATTCCGCGTTACCGTAATTGATGAGGCTGAAATCCGTAGTTTAATATATATTTCACTACTACGGATTTCAGCCTCATCAAAGACGGATACGCGTATGTCAGTTCATCCAAACATAAAC |
| BALF2 (p6927) | shRNA 7323 top  CTAGGTTTATGTTTGGATGAACTGACATACGCGTATCCGTCTTCTCAATCTCATATGTGGTCGGTAGTGAAATATATATTAAACCGACCACATATGAGATTGAGATTACGGTAACGCGG |
|  | shRNA 7323 bot  AATTCCGCGTTACCGTAATCTCAATCTCATATGTGGTCGGTTTAATATATATTTCACTACCGACCACATATGAGATTGAGAAGACGGATACGCGTATGTCAGTTCATCCAAACATAAAC |
|  | shRNA 7324 top  CTAGGTTTATGTTTGGATGAACTGACATACGCGTATCCGTCTTCTTGATCTTGATGTTCCTGGGTAGTGAAATATATATTAAACCCAGGAACATCAAGATCAAGATTACGGTAACGCGG |
|  | shRNA 7324 bot  AATTCCGCGTTACCGTAATCTTGATCTTGATGTTCCTGGGTTTAATATATATTTCACTACCCAGGAACATCAAGATCAAGAAGACGGATACGCGTATGTCAGTTCATCCAAACATAAAC |
|  | shRNA 7325 top  CTAGGTTTATGTTTGGATGAACTGACATACGCGTATCCGTCTTCATAAACTGGACCACTTCGGGTAGTGAAATATATATTAAACCCGAAGTGGTCCAGTTTATGATTACGGTAACGCGG |
|  | shRNA 7325 bot  AATTCCGCGTTACCGTAATCATAAACTGGACCACTTCGGGTTTAATATATATTTCACTACCCGAAGTGGTCCAGTTTATGAAGACGGATACGCGTATGTCAGTTCATCCAAACATAAAC |
| BPLF1 (p6567) | shRNA 6975 top  CTAGGTTTATGTTTGGATGAACTGACATACGCGTATCCGTCTTAATAAACAATTACAGATACAGTAGTGAAATATATATTAAACTGTATCTGTAATTGTTTATTATTACGGTAACGCGG |
|  | shRNA 6975 bot  AATTCCGCGTTACCGTAATAATAAACAATTACAGATACAGTTTAATATATATTTCACTACTGTATCTGTAATTGTTTATTAAGACGGATACGCGTATGTCAGTTCATCCAAACATAAAC |
|  | shRNA 6976 top  CTAGGTTTATGTTTGGATGAACTGACATACGCGTATCCGTCTTTATTAATAAACAATTACAGAGTAGTGAAATATATATTAAACTCTGTAATTGTTTATTAATAATTACGGTAACGCGG |
|  | shRNA 6976 bot  AATTCCGCGTTACCGTAATTATTAATAAACAATTACAGAGTTTAATATATATTTCACTACTCTGTAATTGTTTATTAATAAAGACGGATACGCGTATGTCAGTTCATCCAAACATAAAC |
|  | shRNA 6977 top  CTAGGTTTATGTTTGGATGAACTGACATACGCGTATCCGTCTTCAATAGACAGAAATTGGGTGGTAGTGAAATATATATTAAACCACCCAATTTCTGTCTATTGATTACGGTAACGCGG |
|  | shRNA 6977 bot  AATTCCGCGTTACCGTAATCAATAGACAGAAATTGGGTGGTTTAATATATATTTCACTACCACCCAATTTCTGTCTATTGAAGACGGATACGCGTATGTCAGTTCATCCAAACATAAAC |
| BBRF1 (p6593) | shRNA 6978 for  CTAGGTTTATGTTTGGATGAACTGACATACGCGTATCCGTCTAAAGAATCTTGATAAAACAGGGTAGTGAAATATATATTAAACCCTGTTTTATCAAGATTCTTTTTACGGTAACGCGG |
|  | shRNA 6978 rev  AATTCCGCGTTACCGTAAAAAGAATCTTGATAAAACAGGGTTTAATATATATTTCACTACCCTGTTTTATCAAGATTCTTTAGACGGATACGCGTATGTCAGTTCATCCAAACATAAAC |
|  | shRNA 6979 for  CTAGGTTTATGTTTGGATGAACTGACATACGCGTATCCGTCTTGGCATTCTCAATCATCGAGAGTAGTGAAATATATATTAAACTCTCGATGATTGAGAATGCCATTACGGTAACGCGG |
|  | shRNA 6979 rev  AATTCCGCGTTACCGTAATGGCATTCTCAATCATCGAGAGTTTAATATATATTTCACTACTCTCGATGATTGAGAATGCCAAGACGGATACGCGTATGTCAGTTCATCCAAACATAAAC |
|  | shRNA 6980 for  CTAGGTTTATGTTTGGATGAACTGACATACGCGTATCCGTCTTCATGTTGAACATGACCTCAGGTAGTGAAATATATATTAAACCTGAGGTCATGTTCAACATGATTACGGTAACGCGG |
|  | shRNA 6980 rev  AATTCCGCGTTACCGTAATCATGTTGAACATGACCTCAGGTTTAATATATATTTCACTACCTGAGGTCATGTTCAACATGAAGACGGATACGCGTATGTCAGTTCATCCAAACATAAAC |
| BGLF4 (p6561) | shRNA 6969 top (same as 6966 top)  CTAGGTTTATGTTTGGATGAACTGACATACGCGTATCCGTCTTGATGAAGATGTTGACTGGGAGTAGTGAAATATATATTAAACTCCCAGTCAACATCTTCATCATTACGGTAACGCGG |
|  | shRNA 6969 bot (same as 6966 bot)  AATTCCGCGTTACCGTAATGATGAAGATGTTGACTGGGAGTTTAATATATATTTCACTACTCCCAGTCAACATCTTCATCAAGACGGATACGCGTATGTCAGTTCATCCAAACATAAAC |
|  | shRNA 6970 top  CTAGGTTTATGTTTGGATGAACTGACATACGCGTATCCGTCTAAATCTGATAAATGACCTCTTGTAGTGAAATATATATTAAACAAGAGGTCATTTATCAGATTTTTACGGTAACGCGG |
|  | shRNA 6970 bot  AATTCCGCGTTACCGTAAAAATCTGATAAATGACCTCTTGTTTAATATATATTTCACTACAAGAGGTCATTTATCAGATTTAGACGGATACGCGTATGTCAGTTCATCCAAACATAAAC |
|  | shRNA 6971 top  CTAGGTTTATGTTTGGATGAACTGACATACGCGTATCCGTCTGAAGTAATCAATGACAGTCACGTAGTGAAATATATATTAAACGTGACTGTCATTGATTACTTCTTACGGTAACGCGG |
|  | shRNA 6971 bot  AATTCCGCGTTACCGTAAGAAGTAATCAATGACAGTCACGTTTAATATATATTTCACTACGTGACTGTCATTGATTACTTCAGACGGATACGCGTATGTCAGTTCATCCAAACATAAAC |
| BFRF1 (p6645) | shRNA 6992 top  CTAGGTTTATGTTTGGATGAACTGACATACGCGTATCCGTCTTTATTAATAAAGTGCATACACGTAGTGAAATATATATTAAACGTGTATGCACTTTATTAATAATTACGGTAACGCGG |
|  | shRNA 6992 bot  AATTCCGCGTTACCGTAATTATTAATAAAGTGCATACACGTTTAATATATATTTCACTACGTGTATGCACTTTATTAATAAAGACGGATACGCGTATGTCAGTTCATCCAAACATAAAC |
|  | shRNA 6993 top  CTAGGTTTATGTTTGGATGAACTGACATACGCGTATCCGTCTTGGATATCACAAACACGGGCGGTAGTGAAATATATATTAAACCGCCCGTGTTTGTGATATCCATTACGGTAACGCGG |
|  | shRNA 6993 bot  AATTCCGCGTTACCGTAATGGATATCACAAACACGGGCGGTTTAATATATATTTCACTACCGCCCGTGTTTGTGATATCCAAGACGGATACGCGTATGTCAGTTCATCCAAACATAAAC |
|  | shRNA 6994 top  CTAGGTTTATGTTTGGATGAACTGACATACGCGTATCCGTCTTGAAATTTAGGAAGCAGGGGAGTAGTGAAATATATATTAAACTCCCCTGCTTCCTAAATTTCATTACGGTAACGCGG |
|  | shRNA 6994 bot  AATTCCGCGTTACCGTAATGAAATTTAGGAAGCAGGGGAGTTTAATATATATTTCACTACTCCCCTGCTTCCTAAATTTCAAGACGGATACGCGTATGTCAGTTCATCCAAACATAAAC |
| BSLF1 (p6843) | shRNA 7001 top  CTAGGTTTATGTTTGGATGAACTGACATACGCGTATCCGTCTGATTTGAAAAAATAGACTGGGGTAGTGAAATATATATTAAACCCCAGTCTATTTTTTCAAATCTTACGGTAACGCGG |
|  | shRNA 7001 bot  AATTCCGCGTTACCGTAAGATTTGAAAAAATAGACTGGGGTTTAATATATATTTCACTACCCCAGTCTATTTTTTCAAATCAGACGGATACGCGTATGTCAGTTCATCCAAACATAAAC |
|  | shRNA 7002 top  CTAGGTTTATGTTTGGATGAACTGACATACGCGTATCCGTCTTCTGTGAATAGTACACTGGGGGTAGTGAAATATATATTAAACCCCCAGTGTACTATTCACAGATTACGGTAACGCGG |
|  | shRNA 7002 bot  AATTCCGCGTTACCGTAATCTGTGAATAGTACACTGGGGGTTTAATATATATTTCACTACCCCCAGTGTACTATTCACAGAAGACGGATACGCGTATGTCAGTTCATCCAAACATAAAC |
|  | shRNA 7003 top  CTAGGTTTATGTTTGGATGAACTGACATACGCGTATCCGTCTGTCATTACAAAGTAGTGCCTGGTAGTGAAATATATATTAAACCAGGCACTACTTTGTAATGACTTACGGTAACGCGG |
|  | shRNA 7003 bot  AATTCCGCGTTACCGTAAGTCATTACAAAGTAGTGCCTGGTTTAATATATATTTCACTACCAGGCACTACTTTGTAATGACAGACGGATACGCGTATGTCAGTTCATCCAAACATAAAC |
| BTRF1 (p6845) | shRNA 7004 top  CTAGGTTTATGTTTGGATGAACTGACATACGCGTATCCGTCTGACATTTCTCATAATGGTGCCGTAGTGAAATATATATTAAACGGCACCATTATGAGAAATGTCTTACGGTAACGCGG |
|  | shRNA 7004 bot  AATTCCGCGTTACCGTAAGACATTTCTCATAATGGTGCCGTTTAATATATATTTCACTACGGCACCATTATGAGAAATGTCAGACGGATACGCGTATGTCAGTTCATCCAAACATAAAC |
|  | shRNA 7005 top  CTAGGTTTATGTTTGGATGAACTGACATACGCGTATCCGTCTTTCCTAGAAACTGTGCCATGGGTAGTGAAATATATATTAAACCCATGGCACAGTTTCTAGGAATTACGGTAACGCGG |
|  | shRNA 7005 bot  AATTCCGCGTTACCGTAATTCCTAGAAACTGTGCCATGGGTTTAATATATATTTCACTACCCATGGCACAGTTTCTAGGAAAGACGGATACGCGTATGTCAGTTCATCCAAACATAAAC |
|  | shRNA 7006 top  CTAGGTTTATGTTTGGATGAACTGACATACGCGTATCCGTCTGTGGAAGACAATCTGTCCCGAGTAGTGAAATATATATTAAACTCGGGACAGATTGTCTTCCACTTACGGTAACGCGG |
|  | shRNA 7006 bot  AATTCCGCGTTACCGTAAGTGGAAGACAATCTGTCCCGAGTTTAATATATATTTCACTACTCGGGACAGATTGTCTTCCACAGACGGATACGCGTATGTCAGTTCATCCAAACATAAAC |
| BMRF2 (p6539) | shRNA 6963 for  CTAGGTTTATGTTTGGATGAACTGACATACGCGTATCCGTCTAGGATTTAATGAATGTCACCAGTAGTGAAATATATATTAAACTGGTGACATTCATTAAATCCTTTACGGTAACGCGG |
|  | shRNA 6963 rev  AATTCCGCGTTACCGTAAAGGATTTAATGAATGTCACCAGTTTAATATATATTTCACTACTGGTGACATTCATTAAATCCTAGACGGATACGCGTATGTCAGTTCATCCAAACATAAAC |
|  | shRNA 6964 for  CTAGGTTTATGTTTGGATGAACTGACATACGCGTATCCGTCTTCACAAACTTCTTAAGCTTGTGTAGTGAAATATATATTAAACACAAGCTTAAGAAGTTTGTGATTACGGTAACGCGG |
|  | shRNA 6964 rev  AATTCCGCGTTACCGTAATCACAAACTTCTTAAGCTTGTGTTTAATATATATTTCACTACACAAGCTTAAGAAGTTTGTGAAGACGGATACGCGTATGTCAGTTCATCCAAACATAAAC |
|  | shRNA 6965 for  CTAGGTTTATGTTTGGATGAACTGACATACGCGTATCCGTCTCACAAACTTCTTAAGCTTGTAGTAGTGAAATATATATTAAACTACAAGCTTAAGAAGTTTGTGTTACGGTAACGCGG |
|  | shRNA 6965 rev  AATTCCGCGTTACCGTAACACAAACTTCTTAAGCTTGTAGTTTAATATATATTTCACTACTACAAGCTTAAGAAGTTTGTGAGACGGATACGCGTATGTCAGTTCATCCAAACATAAAC |
| BBRF3 (p6534) | shRNA 6960 for  CTAGGTTTATGTTTGGATGAACTGACATACGCGTATCCGTCTAAAGTAAAAGCTGTTGCCCAGGTAGTGAAATATATATTAAACCTGGGCAACAGCTTTTACTTTTTACGGTAACGCGG |
|  | shRNA 6960 rev  AATTCCGCGTTACCGTAAAAAGTAAAAGCTGTTGCCCAGGTTTAATATATATTTCACTACCTGGGCAACAGCTTTTACTTTAGACGGATACGCGTATGTCAGTTCATCCAAACATAAAC |
|  | shRNA 6961 for  CTAGGTTTATGTTTGGATGAACTGACATACGCGTATCCGTCTAAATAATTTGCAAAGGGCGTGGTAGTGAAATATATATTAAACCACGCCCTTTGCAAATTATTTTTACGGTAACGCGG |
|  | shRNA 6961 rev  AATTCCGCGTTACCGTAAAAATAATTTGCAAAGGGCGTGGTTTAATATATATTTCACTACCACGCCCTTTGCAAATTATTTAGACGGATACGCGTATGTCAGTTCATCCAAACATAAAC |
|  | shRNA 6962 for  CTAGGTTTATGTTTGGATGAACTGACATACGCGTATCCGTCTTTTAAATAATTTGCAAAGGGCGTAGTGAAATATATATTAAACGCCCTTTGCAAATTATTTAAATTACGGTAACGCGG |
|  | shRNA 6962 rev  AATTCCGCGTTACCGTAATTTAAATAATTTGCAAAGGGCGTTTAATATATATTTCACTACGCCCTTTGCAAATTATTTAAAAGACGGATACGCGTATGTCAGTTCATCCAAACATAAAC |
| BNRF1 (p6564) | shRNA 6972 top  CTAGGTTTATGTTTGGATGAACTGACATACGCGTATCCGTCTTCAGTGTATGCATAGTCTGGAGTAGTGAAATATATATTAAACTCCAGACTATGCATACACTGATTACGGTAACGCGG |
|  | shRNA 6972 bot  AATTCCGCGTTACCGTAATCAGTGTATGCATAGTCTGGAGTTTAATATATATTTCACTACTCCAGACTATGCATACACTGAAGACGGATACGCGTATGTCAGTTCATCCAAACATAAAC |
|  | shRNA 6973 top  CTAGGTTTATGTTTGGATGAACTGACATACGCGTATCCGTCTTTGTAAATACAGCACACAGGTGTAGTGAAATATATATTAAACACCTGTGTGCTGTATTTACAATTACGGTAACGCGG |
|  | shRNA 6973 bot  AATTCCGCGTTACCGTAATTGTAAATACAGCACACAGGTGTTTAATATATATTTCACTACACCTGTGTGCTGTATTTACAAAGACGGATACGCGTATGTCAGTTCATCCAAACATAAAC |
|  | shRNA 6974 top  CTAGGTTTATGTTTGGATGAACTGACATACGCGTATCCGTCTGCTATTGCATTAACGAAGGGAGTAGTGAAATATATATTAAACTCCCTTCGTTAATGCAATAGCTTACGGTAACGCGG |
|  | shRNA 6974 bot  AATTCCGCGTTACCGTAAGCTATTGCATTAACGAAGGGAGTTTAATATATATTTCACTACTCCCTTCGTTAATGCAATAGCAGACGGATACGCGTATGTCAGTTCATCCAAACATAAAC |
| BBLF1 (p6557) | shRNA 6966 for  CTAGGTTTATGTTTGGATGAACTGACATACGCGTATCCGTCTTGATGAAGATGTTGACTGGGAGTAGTGAAATATATATTAAACTCCCAGTCAACATCTTCATCATTACGGTAACGCGG |
|  | shRNA 6966 rev  AATTCCGCGTTACCGTAATGATGAAGATGTTGACTGGGAGTTTAATATATATTTCACTACTCCCAGTCAACATCTTCATCAAGACGGATACGCGTATGTCAGTTCATCCAAACATAAAC |
|  | shRNA 6967 for  CTAGGTTTATGTTTGGATGAACTGACATACGCGTATCCGTCTTCATCATTTTCAGAGTCCTCAGTAGTGAAATATATATTAAACTGAGGACTCTGAAAATGATGATTACGGTAACGCGG |
|  | shRNA 6967 rev  AATTCCGCGTTACCGTAATCATCATTTTCAGAGTCCTCAGTTTAATATATATTTCACTACTGAGGACTCTGAAAATGATGAAGACGGATACGCGTATGTCAGTTCATCCAAACATAAAC |
|  | shRNA 6968 for  CTAGGTTTATGTTTGGATGAACTGACATACGCGTATCCGTCTGTCAAAAGTATTGTCTGCGTAGTAGTGAAATATATATTAAACTACGCAGACAATACTTTTGACTTACGGTAACGCGG |
|  | shRNA 6968 rev  AATTCCGCGTTACCGTAAGTCAAAAGTATTGTCTGCGTAGTTTAATATATATTTCACTACTACGCAGACAATACTTTTGACAGACGGATACGCGTATGTCAGTTCATCCAAACATAAAC |
| BRLF1 (p6841) | shRNA 6998 top  CTAGGTTTATGTTTGGATGAACTGACATACGCGTATCCGTCTTTACTATAACTACATTCAGGGGTAGTGAAATATATATTAAACCCCTGAATGTAGTTATAGTAATTACGGTAACGCGG |
|  | shRNA 6998 bot  AATTCCGCGTTACCGTAATTACTATAACTACATTCAGGGGTTTAATATATATTTCACTACCCCTGAATGTAGTTATAGTAAAGACGGATACGCGTATGTCAGTTCATCCAAACATAAAC |
|  | shRNA 6999 top  CTAGGTTTATGTTTGGATGAACTGACATACGCGTATCCGTCTCATCATTTAGAAATGTATCCAGTAGTGAAATATATATTAAACTGGATACATTTCTAAATGATGTTACGGTAACGCGG |
|  | shRNA 6999 bot  AATTCCGCGTTACCGTAACATCATTTAGAAATGTATCCAGTTTAATATATATTTCACTACTGGATACATTTCTAAATGATGAGACGGATACGCGTATGTCAGTTCATCCAAACATAAAC |
|  | shRNA 7000 top  CTAGGTTTATGTTTGGATGAACTGACATACGCGTATCCGTCTACTATAACTACATTCAGGGATGTAGTGAAATATATATTAAACATCCCTGAATGTAGTTATAGTTTACGGTAACGCGG |
|  | shRNA 7000 bot  AATTCCGCGTTACCGTAAACTATAACTACATTCAGGGATGTTTAATATATATTTCACTACATCCCTGAATGTAGTTATAGTAGACGGATACGCGTATGTCAGTTCATCCAAACATAAAC |
| BALF1 (p6632) | shRNA 6986 top  CTAGGTTTATGTTTGGATGAACTGACATACGCGTATCCGTCTTCATTTACAAAGATTTCAGGAGTAGTGAAATATATATTAAACTCCTGAAATCTTTGTAAATGATTACGGTAACGCGG |
|  | shRNA 6986 bot  AATTCCGCGTTACCGTAATCATTTACAAAGATTTCAGGAGTTTAATATATATTTCACTACTCCTGAAATCTTTGTAAATGAAGACGGATACGCGTATGTCAGTTCATCCAAACATAAAC |
|  | shRNA 6987 top  CTAGGTTTATGTTTGGATGAACTGACATACGCGTATCCGTCTATTCATTTACAAAGATTTCAGGTAGTGAAATATATATTAAACCTGAAATCTTTGTAAATGAATTTACGGTAACGCGG |
|  | shRNA 6987 bot  AATTCCGCGTTACCGTAAATTCATTTACAAAGATTTCAGGTTTAATATATATTTCACTACCTGAAATCTTTGTAAATGAATAGACGGATACGCGTATGTCAGTTCATCCAAACATAAAC |
|  | shRNA 6988 top  CTAGGTTTATGTTTGGATGAACTGACATACGCGTATCCGTCTTTACAAAGATTTCAGGAAGTCGTAGTGAAATATATATTAAACGACTTCCTGAAATCTTTGTAATTACGGTAACGCGG |
|  | shRNA 6988 bot  AATTCCGCGTTACCGTAATTACAAAGATTTCAGGAAGTCGTTTAATATATATTTCACTACGACTTCCTGAAATCTTTGTAAAGACGGATACGCGTATGTCAGTTCATCCAAACATAAAC |
| BBRF2 (p6639) | shRNA 6989 top (same as 6978 top)  CTAGGTTTATGTTTGGATGAACTGACATACGCGTATCCGTCTAAAGAATCTTGATAAAACAGGGTAGTGAAATATATATTAAACCCTGTTTTATCAAGATTCTTTTTACGGTAACGCGG |
|  | shRNA 6989 bot (same as 6978 bot)  AATTCCGCGTTACCGTAAAAAGAATCTTGATAAAACAGGGTTTAATATATATTTCACTACCCTGTTTTATCAAGATTCTTTAGACGGATACGCGTATGTCAGTTCATCCAAACATAAAC |
|  | shRNA 6990 top (same as 6979 top)  CTAGGTTTATGTTTGGATGAACTGACATACGCGTATCCGTCTTGGCATTCTCAATCATCGAGAGTAGTGAAATATATATTAAACTCTCGATGATTGAGAATGCCATTACGGTAACGCGG |
|  | shRNA 6990 bot (same as 6979 bot)  AATTCCGCGTTACCGTAATGGCATTCTCAATCATCGAGAGTTTAATATATATTTCACTACTCTCGATGATTGAGAATGCCAAGACGGATACGCGTATGTCAGTTCATCCAAACATAAAC |
|  | shRNA 6991 top  CTAGGTTTATGTTTGGATGAACTGACATACGCGTATCCGTCTTGGTCAATAAAGAATCTTGATGTAGTGAAATATATATTAAACATCAAGATTCTTTATTGACCATTACGGTAACGCGG |
|  | shRNA 6991 bot  AATTCCGCGTTACCGTAATGGTCAATAAAGAATCTTGATGTTTAATATATATTTCACTACATCAAGATTCTTTATTGACCAAGACGGATACGCGTATGTCAGTTCATCCAAACATAAAC |
| BMLF1 (p6838) | shRNA 6995 top  CTAGGTTTATGTTTGGATGAACTGACATACGCGTATCCGTCTTGTAATTCTTGATGTAGTGGCGTAGTGAAATATATATTAAACGCCACTACATCAAGAATTACATTACGGTAACGCGG |
|  | shRNA 6995 bot  AATTCCGCGTTACCGTAATGTAATTCTTGATGTAGTGGCGTTTAATATATATTTCACTACGCCACTACATCAAGAATTACAAGACGGATACGCGTATGTCAGTTCATCCAAACATAAAC |
|  | shRNA 6996 top  CTAGGTTTATGTTTGGATGAACTGACATACGCGTATCCGTCTTTATTGATTTAATCCAGGAACGTAGTGAAATATATATTAAACGTTCCTGGATTAAATCAATAATTACGGTAACGCGG |
|  | shRNA 6996 bot  AATTCCGCGTTACCGTAATTATTGATTTAATCCAGGAACGTTTAATATATATTTCACTACGTTCCTGGATTAAATCAATAAAGACGGATACGCGTATGTCAGTTCATCCAAACATAAAC |
|  | shRNA 6997 top  CTAGGTTTATGTTTGGATGAACTGACATACGCGTATCCGTCTTCACAAAGTTGTAGTCTCGCGGTAGTGAAATATATATTAAACCGCGAGACTACAACTTTGTGATTACGGTAACGCGG |
|  | shRNA 6997 bot  AATTCCGCGTTACCGTAATCACAAAGTTGTAGTCTCGCGGTTTAATATATATTTCACTACCGCGAGACTACAACTTTGTGAAGACGGATACGCGTATGTCAGTTCATCCAAACATAAAC |
|  | Antisense/target: red-colored (top, for) and brown-colored (bot, rev)  Sense/passenger: green-colored (top, for) and blue-colored (bot, rev) |
