## Supplementary Table S3 for "Systematic analysis of Epstein-Barr virus genes and their individual contribution to virus production and composition"

**Table S3. Synthetic pairs of oligonucleotides used for the construction of luciferase reporter plasmids**

| Target gene (plasmid #) | Oligonucleotide |
| --- | --- |
| BALF4 (p7204) | 7204 for (sh6515-2, BALF4)  TCGAGATGGTGTTTAAAGACAACATTAAATTTGCAGACCTTCATCTCACTAAAAATTGAAGCCATTACATATTTTATAAGC |
|  | 7204 rev (sh6515-2, BALF4)  GGCCGCTTATAAAATATGTAATGGCTTCAATTTTTAGTGAGATGAAGGTCTGCAAATTTAATGTTGTCTTTAAACACCATC |
| BKRF4 (p7329) | 7329 top (sh6562, BKRF4)  TCGAGCCCAATACATACAATAAACACAAATTAAGTCCCAATACATACAATAAAAATTTCAAGTCCCAATACATACAATAGC |
|  | 7329 bot (sh6562, BKRF4)  GGCCGCTATTGTATGTATTGGGACTTGAAATTTTTATTGTATGTATTGGGACTTAATTTGTGTTTATTGTATGTATTGGGC |
| BVLF1 (p7330) | 7330 top (sh6846, BVLF1)  TCGAGCCCCGTGCGATGAGTTTATTTAAATTTCCTCCTGAAGATGCTAAAGAAAATTTGACCCTAAGTGTTAACTTTTAGC |
|  | 7330 bot (sh6846, BVLF1)  GGCCGCTAAAAGTTAACACTTAGGGTCAAATTTTCTTTAGCATCTTCAGGAGGAAATTTAAATAAACTCATCGCACGGGGC |
| BNLF2a (p7331) | 7331 top (sh6839, BNLF2a)  TCGAGGACTTGTGATGCAATAAATAAAAATTTGACTTGTGATGCAATAAATAAAATTATGTGACTTGTGATGCAATAAAGC |
|  | 7331 bot (sh6839, BNLF2a)  GGCCGCTTTATTGCATCACAAGTCACATAATTTTATTTATTGCATCACAAGTCAAATTTTTATTTATTGCATCACAAGTCC |
| BXRF1 (p7332) | 7332 top (sh6848, BXRF1)  TCGAGTCGCGGACAATAAGATTTACAAAATTGGTGTTATATTGTAGAATTTAAAATTACACCCTCTTTGAGATTATCAAGC |
|  | 7332 bot (sh6848, BXRF1)  GGCCGCTTGATAATCTCAAAGAGGGTGTAATTTTAAATTCTACAATATAACACCAATTTTGTAAATCTTATTGTCCGCGAC |
| BFLF2 (p7333) | 7333 top (sh6644, BFLF2)  TCGAGCCCAGCTCATTTTGGAAAATAAAATTCCAGCTCATTTTGGAAAATAAAAATTGACCTGGTACAGTCCTATCAAAGC |
|  | 7333 bot (sh6644, BFLF2)  GGCCGCTTTGATAGGACTGTACCAGGTCAATTTTTATTTTCCAAAATGAGCTGGAATTTTATTTTCCAAAATGAGCTGGGC |
| BMRF1 (p7334) | 7334 top (sh5106, BMRF1)  TCGAGTGGTGACATTCATTAAATCCTAAATTGTTGGATCTTAGTGTTATTTTAAATTTCACTATGGAATATGATGATAAGC |
|  | 7334 bot (sh5106, BMRF1)  GGCCGCTTATCATCATATTCCATAGTGAAATTTAAAATAACACTAAGATCCAACAATTTAGGATTTAATGAATGTCACCAC |
| BHLF1 (p7335) | 7335 top (sh7080, BHLF1)  TCGAGCACCTATCAATAAACTGTTTAAAATTACACTACTCACCTACATGTCAAAATTGTACACTACACTCTAAAAGTAAGC |
|  | 7335 bot (sh7080, BHLF1)  GGCCGCTTACTTTTAGAGTGTAGTGTACAATTTTGACATGTAGGTGAGTAGTGTAATTTTAAACAGTTTATTGATAGGTGC |
| BXLF1 (p7337) | 7337 top (sh6571, BXLF1)  TCGAGACAGCATCTTTAGTGTACTTAAAATTACCGCTACAAGTTAATTTACAAAATTTGGGCGCGAGTTTAGTAAGTTAGC |
|  | 7337 bot (sh6571, BXLF1)  GGCCGCTAACTTACTAAACTCGCGCCCAAATTTTGTAAATTAACTTGTAGCGGTAATTTTAAGTACACTAAAGATGCTGTC |
| BVRF2 (p7267) | 7267 for (sh6847, BVRF2)  TCGAGTCCCAGCCGAGTCGTATTTAAAAATTCACAGGTAAAAGTTAACACTTAAATTTACGGATTTCAGCCTCATCAAAGC |
|  | 7267 rev (sh6847, BVRF2)  GGCCGCTTTGATGAGGCTGAAATCCGTAAATTTAAGTGTTAACTTTTACCTGTGAATTTTTAAATACGACTCGGCTGGGAC |
| BALF2 (p7338) | 7338 top (sh6927, BALF2)  TCGAGCGACCACATATGAGATTGAGAAAATTCCAGGAACATCAAGATCAAGAAAATTCCGAAGTGGTCCAGTTTATGAAGC |
|  | 7338 bot (sh6927, BALF2)  GGCCGCTTCATAAACTGGACCACTTCGGAATTTTCTTGATCTTGATGTTCCTGGAATTTTCTCAATCTCATATGTGGTCGC |
| BPLF1 (p7211) | 7211 for (sh6567-2, BPLF1)  TCGAGTGTATCTGTAATTGTTTATTAAAATTTCTGTAATTGTTTATTAATAAAAATTCACCCAATTTCTGTCTATTGAAGC |
|  | 7211 rev (sh6567-2, BPLF1)  GGCCGCTTCAATAGACAGAAATTGGGTGAATTTTTATTAATAAACAATTACAGAAATTTTAATAAACAATTACAGATACAC |
| BBRF1 (p7212) | 7212 for (sh6593-4, BRRF1)  TCGAGCCTGTTTTATCAAGATTCTTTAAATTTCTCGATGATTGAGAATGCCAAAATTCTGAGGTCATGTTCAACATGAAGC |
|  | 7212 rev (sh6593-4, BRRF1)  GGCCGCTTCATGTTGAACATGACCTCAGAATTTTGGCATTCTCAATCATCGAGAAATTTAAAGAATCTTGATAAAACAGGC |
| BGLF4 (p7209) | 7209 for (sh6561-2, BGLF4)  TCGAGTCCCAGTCAACATCTTCATCAAAATTAAGAGGTCATTTATCAGATTTAAATTGTGACTGTCATTGATTACTTCAGC |
|  | 7209 rev (sh6561-2, BGLF4)  GGCCGCTGAAGTAATCAATGACAGTCACAATTTAAATCTGATAAATGACCTCTTAATTTTGATGAAGATGTTGACTGGGAC |
| BFRF1 (p7215) | 7215 for (sh6645-2, BFRF1)  TCGAGGTGTATGCACTTTATTAATAAAAATTCGCCCGTGTTTGTGATATCCAAAATTTCCCCTGCTTCCTAAATTTCAAGC |
|  | 7215 rev (sh6645-2, BFRF1)  GGCCGCTTGAAATTTAGGAAGCAGGGGAAATTTTGGATATCACAAACACGGGCGAATTTTTATTAATAAAGTGCATACACC |
| BSLF1 (p7218) | 7218 for (sh6843-2, BSLF1)  TCGAGCCCAGTCTATTTTTTCAAATCAAATTCCCCAGTGTACTATTCACAGAAAATTCAGGCACTACTTTGTAATGACAGC |
|  | 7218 rev (sh6843-2, BSLF1)  GGCCGCTGTCATTACAAAGTAGTGCCTGAATTTTCTGTGAATAGTACACTGGGGAATTTGATTTGAAAAAATAGACTGGGC |
| BTRF1 (p7219) | 7219 for (sh6845-2, BTRF1)  TCGAGGGCACCATTATGAGAAATGTCAAATTCCATGGCACAGTTTCTAGGAAAAATTTCGGGACAGATTGTCTTCCACAGC |
|  | 7219 rev (sh6845-2, BTRF1)  GGCCGCTGTGGAAGACAATCTGTCCCGAAATTTTTCCTAGAAACTGTGCCATGGAATTTGACATTTCTCATAATGGTGCCC |
| BMRF2 (p7207) | 7207 for (sh6539-2, BMRF2)  TCGAGTGGTGACATTCATTAAATCCTAAATTACAAGCTTAAGAAGTTTGTGAAAATTTACAAGCTTAAGAAGTTTGTGAGC |
|  | 7207 rev (sh6539-2, BMRF2)  GGCCGCTCACAAACTTCTTAAGCTTGTAAATTTTCACAAACTTCTTAAGCTTGTAATTTAGGATTTAATGAATGTCACCAC |
| BBRF3 (p7206) | 7206 for (sh6534-4, BRRF3)  TCGAGCTGGGCAACAGCTTTTACTTTAAATTCACGCCCTTTGCAAATTATTTAAATTGCCCTTTGCAAATTATTTAAAAGC |
|  | 7206 rev (sh6534-4, BRRF3)  GGCCGCTTTTAAATAATTTGCAAAGGGCAATTTAAATAATTTGCAAAGGGCGTGAATTTAAAGTAAAAGCTGTTGCCCAGC |
| BNRF1 (p7210) | 7210 for (sh6564-2, BNRF1)  TCGAGTCCAGACTATGCATACACTGAAAATTACCTGTGTGCTGTATTTACAAAAATTTCCCTTCGTTAATGCAATAGCAGC |
|  | 7210 rev (sh6564-2, BNRF1)  GGCCGCTGCTATTGCATTAACGAAGGGAAATTTTTGTAAATACAGCACACAGGTAATTTTCAGTGTATGCATAGTCTGGAC |
| BBLF1 (p7208) | 7208 for (sh6557-2, BBLF1)  TCGAGTCCCAGTCAACATCTTCATCAAAATTTGAGGACTCTGAAAATGATGAAAATTTACGCAGACAATACTTTTGACAGC |
|  | 7208 rev (sh6557-2, BBLF1)  GGCCGCTGTCAAAAGTATTGTCTGCGTAAATTTTCATCATTTTCAGAGTCCTCAAATTTTGATGAAGATGTTGACTGGGAC |
| BRLF1 (p7217) | 7217 for (sh6841-2, BRLF1)  TCGAGCCCTGAATGTAGTTATAGTAAAAATTTGGATACATTTCTAAATGATGAAATTATCCCTGAATGTAGTTATAGTAGC |
|  | 7217 rev (sh6841-2, BRLF1)  GGCCGCTACTATAACTACATTCAGGGATAATTTCATCATTTAGAAATGTATCCAAATTTTTACTATAACTACATTCAGGGC |
| BALF1 (p7213) | 7213 for (sh6632-2, BALF1)  TCGAGTCCTGAAATCTTTGTAAATGAAAATTCTGAAATCTTTGTAAATGAATAAATTGACTTCCTGAAATCTTTGTAAAGC |
|  | 7213 rev (sh6632-2, BALF1)  GGCCGCTTTACAAAGATTTCAGGAAGTCAATTTATTCATTTACAAAGATTTCAGAATTTTCATTTACAAAGATTTCAGGAC |
| BBRF2 (p7214) | 7214 for (sh6639-2, BBRF2)  TCGAGCCTGTTTTATCAAGATTCTTTAAATTTCTCGATGATTGAGAATGCCAAAATTATCAAGATTCTTTATTGACCAAGC |
|  | 7214 rev (sh6639-2, BBRF2)  GGCCGCTTGGTCAATAAAGAATCTTGATAATTTTGGCATTCTCAATCATCGAGAAATTTAAAGAATCTTGATAAAACAGGC |
| BMLF1 (p7216) | 7216 for (sh6838-2, BMLF1)  TCGAGGCCACTACATCAAGAATTACAAAATTGTTCCTGGATTAAATCAATAAAAATTCGCGAGACTACAACTTTGTGAAGC |
|  | 7216 rev (sh6838-2, BMLF1)  GGCCGCTTCACAAAGTTGTAGTCTCGCGAATTTTTATTGATTTAATCCAGGAACAATTTTGTAATTCTTGATGTAGTGGCC |
|  | Sense/target: three individual red-colored sequences  Antisense: three individual brown-colored sequences |
